# Circadian desynchronization desensitizes insulin-producing cells to cytokine-mediated transcriptomic remodeling and cell death: a novel β-cell anti-apoptotic response to inflammation

**DOI:** 10.1101/2024.04.12.589193

**Authors:** Phillip Alexander Keller Andersen, Rasmus Henrik Reeh, Isabel Sanders, Emilie Bender Overlund, Thomas Mandrup-Poulsen

## Abstract

Perturbation of the circadian clock is a risk factor for metabolic diseases. β-Cell specific clock disruption causes glucose intolerance in mice, associated with oxidative stress and secretory failure in β-cells. Proinflammatory cytokines alter the expression of core-clock machinery in human and rodent β-cells, but the molecular mechanisms and consequences for cell viability are unclear. We hypothesized that cytokine-mediated clock perturbation in β-cells is NF-κB driven, concomitant with cytokine-induced apoptosis, and depends on the cellular synchronization status. Cytokine-mediated changes of core-clock mRNA expression observed in non-synchronized INS-1 cells were potentiated in synchronized cells. These transcriptional changes differentially translated into alterations in core-clock protein levels. Interestingly, synchronization also sensitized INS-1 cells to cytokine-mediated cytotoxicity, associated with potentiation in the expression of inducible (ind) proteasomal catalytic subunits, ER stress markers, NF-κB activity, and activation of the intrinsic apoptotic pathway. Small-molecule NF- κB inhibition abrogated cytokine-mediated regulation of clock gene expression in both synchronized and non-synchronized INS-1 cells and reversed cytokine-mediated alterations in circadian parameters in INS-1 reporter cells at non-cytotoxic concentrations. However, at cytotoxic cytokine concentrations, NF-κB inhibition caused a loss of circadian rhythmicity while still reducing the cytotoxic effects of cytokines, indicating a differential effect of NF-κB signaling in controlling β-cell viability and clock regulation. We propose that in synchronized cells, the proinflammatory transcriptional activity of NF-κB is enhanced by interaction with clock transcription factors, as has been suggested for the clock activator Brain and muscle Arnt-like protein-1 (Bmal1). Thus, desynchronization provides a novel anti-apoptotic defense mechanism in response to cytokine assault, similar to that provided by β-cell phenotypic de-differentiation.

## Introduction

The circadian system, highly conserved in evolution, enables light-sensitive organisms to anticipate, prepare for, and respond to regularly recurring physiological needs during a solar day [1]. To uphold circadian regulation at the systemic organismal level, almost all cells harbor intrinsic molecular clocks, that operate via interlocked transcriptional-translational feedback loops [1]. Individual cellular molecular clocks are synchronized through multiple entrainment signals (*Zeitgebers*) [2] comprising light [3], temperature [4], nutrition [5], and physical activity [6] that are integrated by the central clock in the suprachiasmatic nuclei (SCN) to convey synchronizing neuro-humoral signals to the peripheral oscillators.

Coordinated circadian oscillations are required for normal tissue functions, and circadian misalignment is a risk factor for common chronic diseases, e.g. neoplastic, cardio-metabolic, and psychiatric disorders [7]. Indeed, circadian disruption and misalignment confer β-cell dysfunction and insulin resistance [8–11], while global or β-cell specific knock-out of core molecular clock activators results in β-cell secretory failure and diabetes [12, 13].

Inflammation is a common pathogenetic factor in both type 1 and type 2 diabetes, and there is clinical proof-of-concept that proinflammatory cytokines contribute to β-cell failure in these diseases [14–16]. The molecular mechanisms underlying cytokine-mediated β-cell failure and destruction are incompletely understood but involve Nuclear factor of kappa light polypeptide gene enhancer in B-cells (NF-κB) and Mitogen activated protein kinase (MAPK) activation, and nitroxidative, endoplasmic reticulum (ER) and mitochondrial stress that eventually trigger the intrinsic apoptosis program [17, 18].

NF-κB activation induces transcriptional reprogramming in β-cells, with extensive alterations of the expressional profiles of hundreds of both defensive and deleterious pathways. Many of the cytokine-induced β-cell transcriptomic changes ultimately causing secretory dysfunction and culminating in apoptosis are NO and thereby NF-κB dependent [18–21]. Interestingly, in addition to its role in inflammation, NF-κB signaling is required for a functional molecular clock [22, 23], indicating a context-dependent interplay between physiological and pathophysiological actions of NF-κB.

We and others [24, 25] have recently found that proinflammatory cytokines reconfigure the expression pattern of core-clock genes, while also changing rhythmic parameters in islets harboring a Period2 promotor-driven Luciferase (*Per2-Luc*). We found that the proinflammatory cytokines Interleukin-1β (IL-1β) and Interferon-γ (IFN-γ) increased period length in both human and murine reporter islets [24]. Further, we observed that Brain and muscle arnt-like 1 (*Bmal1*), Circadian locomotor output cycles kaput (*Clock*), Nuclear receptor subfamily 1 group D member 1 (*Nr1d1* or *Rev-erbα*), *Per1, Per2* and Cryptochrome 2 (*Cry2*) were upregulated, while *Cry1* was downregulated by these cytokines in the rat insulin-producing cell line, INS-1. Interestingly, these apparently uncoordinated expressional changes were reversed by inhibition of inducible nitric oxide synthase (iNOS), histone deacetylase 3 (HDAC3), and the immune/intermediate proteasome. All these pathways are involved in inflammatory β-cell stress, and regulated by NF-κB [14, 18, 24, 26]. However, a direct demonstration that NF-κB signaling conveys proinflammatory stress-mediated clock gene alterations and period perturbation in β-cells is lacking. Furthermore, a main limitation in [24] was that gene expression levels were only determined in a non-synchronized model system without parallel documentation of β-cell viability under these conditions.

Cultured cells can synchronize *in vitro* using different Z*eitgebers*, such as serum shock, dexamethasone, forskolin pulsing, and many more [27, 28]. There is emerging evidence that synchronized cell models are more sensitive to chemical toxins compared to non-synchronized cells [29]. Accordingly, we hypothesized that cytokine-mediated clock perturbation is NF-κB driven, as is cytokine-induced apoptosis but depends on the cellular synchronization status and the intensity of inflammatory stress. We demonstrate that cytokine-mediated β-cell clock perturbation is NF-κB dependent, and that synchronization of cellular rhythms potentiated proinflammatory stress-mediated expressional alterations of selected core-clock genes and proteins, while also potentiating ER stress and associated cytotoxicity in a context- and NF-κB-dependent manner. We suggest that desynchronization provides a novel anti-apoptotic defense in response to cytokine assault, similar to that provided by β-cell phenotypic de-differentiation [30–32].

## Materials and methods

### Cell culture

The rat INS-1 insulinoma cell line was kindly provided by C. Wollheim and P. Maechler, University Medical Centre, Geneva, Switzerland. Cells were cultured in RPMI-1640 with GlutaMAX™, and supplemented with 10% FBS, 1% penicillin/streptomycin, HEPES, sodium bicarbonate, and 50μM β-mercaptoethanol (culture medium) (Life Technologies, Carlsbad, CA, USA). The EndoC-βH3 cell line was purchased from Human Cell Design (Toulouse, France) and cultured in medium as described in [33], but without the addition of puromycin. The cells were exposed to proinflammatory cytokines, inhibitors, and conditions in the concentrations and for the time periods specified in figure legends. For functional assays using absorbance, luminescence, or fluorescence, culture medium was used, but without phenol red.

### RT-qPCR

Cells were seeded in a 6 well-plate (0.5×10^6^ INS-1 cells/ well) for 3 days. RNA was extracted using NucleoSpin® RNA kit (Macherey-Nagel, Bethlehem, PA, USA). RNA quality and quantity were assessed using Nanodrop-2000 (Thermo Fisher Scientific, Waltham, MA, USA).

cDNA was synthesized from 500 ng RNA using the iScript™-cDNA Kit (Bio-Rad Laboratories, Hercules, CA, USA). Real-time quantitative PCR was carried out using mRNA-specific primers (Table S1), designed in-lab and synthesized by TAG Copenhagen (Copenhagen, Denmark), and SYBR^TM^ Green PCR Master Mix (Applied Biosystems, Waltham, MA, USA). Amplification and real-time monitoring were carried out using QuantStudio5, and data was analyzed using QuantStudio™ Design & Analysis Software (Applied Biosystems). All reactions were carried out following the manufacturer’s protocols. mRNA expression was visualized either as normalized expression using the 2^-ΔΔCt^ method (ΔCt (exposed sample) – ΔCt (control sample) or as relative expression using the 2^-ΔCt^ method, where ΔCt values are defined as Ct_target gene_-Ct_reference gene_. Statistical differences in relative mRNA expression were determined using log2 transformed data. The stability of the reference mRNA was evaluated based on raw Ct values, using the most stable mRNA for normalization in the given experimental setup, as given in the figure legends. If two or more reference genes provided the least variation in Ct values, the geometric mean of the genes was used.

### Western blotting

Cells were lysed in lysis buffer consisting of 100 mM Tris (pH 8.0), 30 mM NaCl, 10 mM KCl, 10 mM MgCl2, 2% IGEPAL, 20 mM iodoacetamide and Roche Protease inhibitor tablets (Life Technologies). Protein concentrations were measured by Bradford assay using Bio-Rad Protein Assay Dye Reagent (Bio-Rad Laboratories). Indicated amounts of protein were loaded on Nu-Page 4-12% bis-tris gels (Thermo Fisher Scientific), separated by SDS-PAGE, and transferred to PVDF membranes via the iBLOT2 system (Thermo Fisher Scientific). Total protein on the blots was detected using TotalStain Q –PVDF Fluorescent Total Protein Staining Kit (Azure Biosystems, Ohio, USA), and captured using the Azure®Saphire Biomolecular Imager (Azure Biosystems). Membranes were blocked using 5% skim-milk powder (MilliporeSigma, Massachusetts, USA) for one hour, following incubation with primary antibodies (Table S2) diluted in TBST, containing 2% BSA and 0.27% sodium azide or in azure blocking buffer with sodium azide, overnight. Membranes were incubated with the appropriate secondary antibodies (dilution 1:10,000) for two hours. The final blots were developed by means of chemiluminescence using Radiance Q Chemiluminescent Substrate (Azure Biosystems). Image Lab 6.1 (Bio-Rad Laboratories) and AzureSpot Pro (Azure Biosystems) were used for densitometric analysis of the Western Blots. Antibodies and dilutions are annotated in Table S2. Quantified values are normalized using a build-in normalization method in Graphpad Prism 10 (GraphPad Software, Boston, MA, USA), in which 0 is set as 0%, while 100% is determined as the mean of the biological replicates within an experimental condition, of either 8 or 36 hours, depending on which value is greater.

### Lentiviral transduction of INS-1 cells

Lentiviral particles were produced by transfecting 293T cells in 10 cm dishes with helper plasmids Pax2 and MD.2g together with either Period2 promotor-driven Luciferase (Per2-Luc) (a gift from Prof. Louis Ptácek, University of California San Francisco) or GFP lentivectors by calcium phosphate transfection. Virus supernatant was collected after 48 hours and concentrated 20X in Amicon filters with a 100,000 kDa cut-off. INS-1 cells were transduced with either Per2-Luc or GFP lentiviral particles over 48 hours, and approximate transduction efficiency was assessed by GFP transduction in parallel cultures. Per2-luc transduced cells corresponding to a GFP expression of ∼65% (one virus integration per genome) were selected with blasticidin.

### *In vitro* synchronization

INS-1 and EndoC-βH3 cells were synchronized using a one-hour 10 µM forskolin pulse (Sigma-Aldrich, St. Louis, Missouri, USA), followed by washing with 37°C medium, and incubation in medium with the exposures, as in [24, 28]. When cells were exposed to experimental conditions, such as cytokines, while also subjected to *in vitro* synchronization, cells were co-incubated with the experimental condition (e.g. cytokines) and forskolin in the one-hour pulse. At the end of the pulse, cells were washed, and then the experimental conditions were added for the remaining duration of the experiment.

### Circadian bioluminescence recording

INS-1 Per2-Luc cells were incubated for 24 hours in a white-walled and -bottomed 96-well plate (15,000 cells/well). After exposures and forskolin pulse, the cells were incubated in culture medium without phenol red with 0.1 mM D-luciferin (VivoGlo™ Luciferin, Promega, Madison, WI, USA) (experimental medium). Bioluminescence was recorded using the CLARIOstar Plus, with an atmospheric control unit (ACU) (BMG LABTECH, Ortenberg, Germany) to allow for continuous 96-hour monitoring at 10-minute intervals at 37°C and 5% CO_2_. Raw traces were detrended using a 24-hour window moving average as in [9]. The functional window defined below was used to determine rhythmic parameters by the mFourFit method carried out using BioDare2 [34]. At low concentrations of IL-1β (≤15 pg/mL), the functional window was set from 13.5 to 85.5 hours of the recording. At high IL-1β (≥30 pg/mL), the functional window was set from 30 to 85.5 hours of the recording, due to the noisy initial 30-hour period preventing rhythmic analysis, as illustrated in the detrended bioluminescence recordings.

### Viability

INS-1 cells were seeded at 30,000 cells per well in a clear 96-well plate for 3 days, followed by exposures as described in the figure legends. Mitochondrial function was determined using AlamarBlue™ HS Cell Viability Reagent (Invitrogen, Waltham, MA, USA) as a proxy for cell viability, following the manufacturer’s protocol, and absorbance or fluorescence recorded on a SpectraMax® i3x (Molecular Devices, San Jose, CA, USA).

### Cytotoxicity

Total accumulated cytotoxicity (an integrated readout for apoptosis and late apoptotic necrosis) was monitored in real-time using the Cell-tox assay (Promega) according to the manufacturer’s protocol. Cells were seeded in either a black-walled plate, if the toxicity assay was run alone, or in a white-walled plate if multiplexed with the circadian bioluminescence assay. Cells were incubated and monitored in real-time at 10-minute intervals for 72-96 hours using the CLARIOstar Plus, with ACU (BMG LABTECH) at 37°C and 5% CO2.

### Caspase assay

Caspase activity was measured following the manufacturer’s instructions using the Caspase-Glo™ 3/7, 8, and 9 Assays (Promega). Ten thousand cells per well were seeded in a white-walled 96-well plate for 48 hours, followed by exposures for 24 hours. Luminescence was measured using a SpectraMax® i3x (Molecular Devices) after a 30-minute incubation with the assay reagents.

### Bioinformatics

Proteasomal and immunoproteasomal cleavage sites of selected clock proteins were predicted using the “Improved Proteasome Cleavage Prediction Server” (iPCPS) (http://imed.med.ucm.es/Tools/pcps/).

### Statistics

Data were analyzed using GraphPad Prism 9/10 as specified in the figure legends. Data are presented as means ± SEMs. Significance levels are annotated as follows: * = p-value < 0.05, ** = p-value < 0.01, *** = p-value < 0.001, **** = p-value < 0.0001.

## Results

### Proinflammatory cytokines introduce uncoordinated alterations in clock protein expression

As we have previously [24] investigated the effect of proinflammatory cytokines on core-clock gene expression to assess the transcriptional response to inflammatory stress, we first wished to assess the expression levels of selected core-clock proteins to evaluate the actual levels of transcriptional repressors and activators. INS-1 cells were exposed to a combination of the proinflammatory cytokines IL-1β and IFN-γ for 8-36 hours and harvested at 4-hour intervals to allow for analysis of both cytokine-mediated and temporal effects. Protein levels of REV-ERBα, PER2, CRY2, BMAL1, and CLOCK were determined (FIG. S1). These proteins were selected based on the effect sizes of the cytokine combination on their corresponding mRNA levels as reported in [24]. Cytokines upregulated both CLOCK, REV-ERBα, and PER2, and for CLOCK with a significant impact of both cytokine exposure and time dependency, and a trend towards an interaction between these in two-way ANOVA.

Thus, IL-1β and IFN-γ upregulate the expression levels of a subset of the selected core-clock proteins.

### Synchronization potentiates the cytokine-mediated alterations in expression of core-clock mRNAs and proteins, and of inducible proteasome transcripts

To investigate proinflammatory stress-mediated changes in core-clock gene mRNA and protein levels in a relevant circadian model, INS-1 cells harboring *Per2* promotor driven luciferase (*Per2-Luc*), as a readout of integrated circadian rhythmicity, were synchronized *in vitro* by an one-hour 10 µM forskolin pulse, which is a known synchronization stimulus, preferentially used in the synchronization of β-cells [13, 35, 36]. Additionally, cells were subjected to medium change alone, or direct administration of Luciferin (no media change), to control for cell handling. Forskolin induced an oscillatory rhythm distinct from that observed after cell handling (FIG. S2). The forskolin-induced rhythmic oscillations of circadian bioluminescence were concordant with endogenous in core-clock mRNA expression that showed anti-phasic rhythmic profiles between repressors and activators, also observed to a lesser extent at protein levels (FIG. S3-S4). In the following, “synchronized” will refer to cells exposed to a forskolin pulse, while “non-synchronized” will refer to cells not exposed to a forskolin pulse, even if they have some inherent rhythmicity.

Next, synchronized INS-1 cells were exposed to IL-1β and IFN-γ, and the expression levels of the selected clock gene mRNAs (FIG. 1) and proteins (FIG. 2) were investigated in a time-dependent manner. The cytokine-induced mRNA expression of *Rev-erbα*, *Per2,* and *Bmal1* was potentiated by synchronization (FIG. 1), particularly notable at the peaks for *Bmal1* (a 48% increase) and *Rev-erbα* (a 46 % increase). Cytokine-induced *Cry2* expression was decreased, while *Clock* expression was not changed by synchronization.

**Figure 1.**
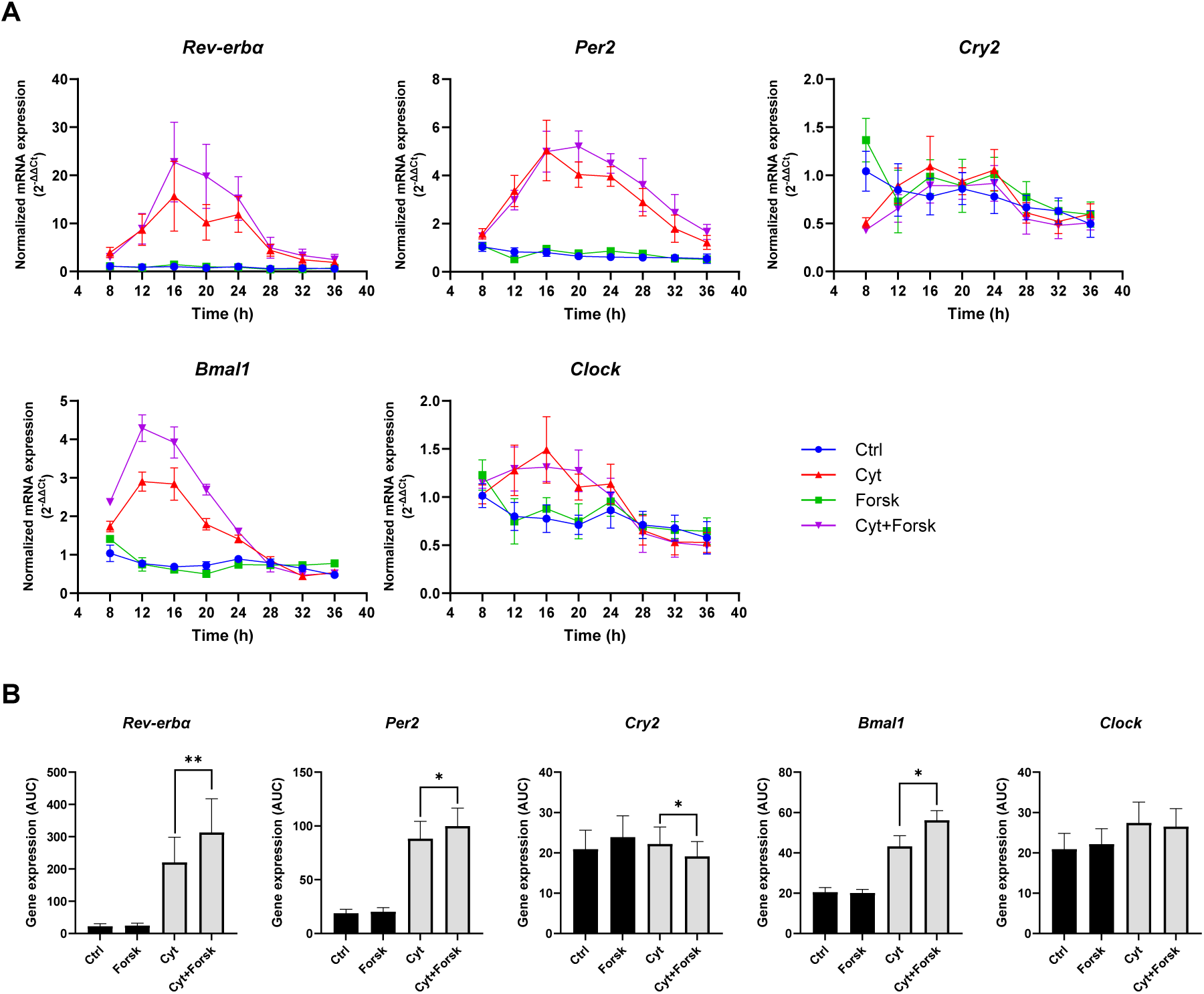
Synchronization potentiates the proinflammatory cytokine-mediated alterations in core-clock gene expression. INS-1 cells were cultured in the presence (Cyt) or absence (Ctrl) of 150 pg/ml of mIL-1β and 0.1 ng/ml rIFN-γ. Additionally, cells were synchronized with a 1-hour forskolin pulse in the absence (Forsk) or presence (Cyt+Forsk) of the same cytokine concentrations, as described in material and methods. Cells were exposed for between 8 and 36 hours, with cell harvest at 4-hour intervals. Normalized mRNA expression was calculated using *Hprt1* as a reference gene. **(A)** Time-dependent normalized mRNA expression. **(B)** AUC of the curves in (A). Data are presented as means ± SEM (N=4). Statistics is two-tailed paired Student’s t-test.

**Figure 2.**
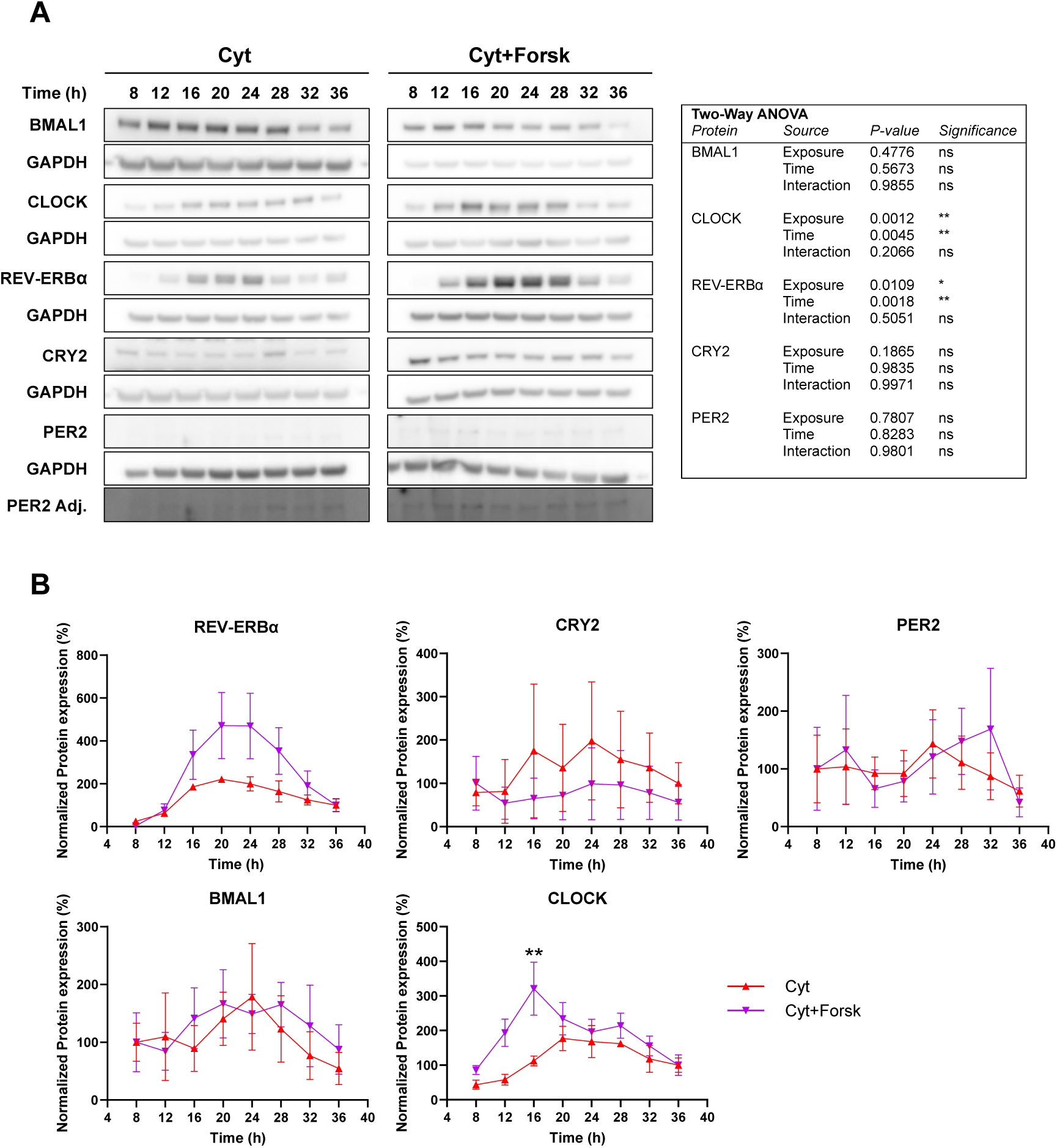
Synchronization potentiates the proinflammatory cytokine-mediated increase in CLOCK and REV-ERBα expression. INS-1 cells were exposed as described in FIG. 1, only using cytokine-exposed cells in a non-synchronized (Cyt) or synchronized (Cyt+Forsk) system. **(A)** Samples were analyzed by SDS-PAGE and Western blotting with GAPDH as the internal loading control. PER2 Adj. is the PER2 band gamma adjusted to allow for illustration of the bands, while the unadjusted PER2 bands were used for quantification **(B)** Quantification and normalization of the protein expression levels. Data are presented as means ± SEM (N=2-4). Statistics are two-way ANOVA with Šídák’s corrected multiple comparisons tests. A summary of the statistical tests is presented in the table.

Cytokine-induced REV-ERBα and CLOCK proteins were significantly increased by synchronization, with the time dependency also being affected (FIG. 3). However, the activator protein BMAL1 and the repressor proteins PER2 and CRY2 were unchanged after synchronization.

**Figure 3.**
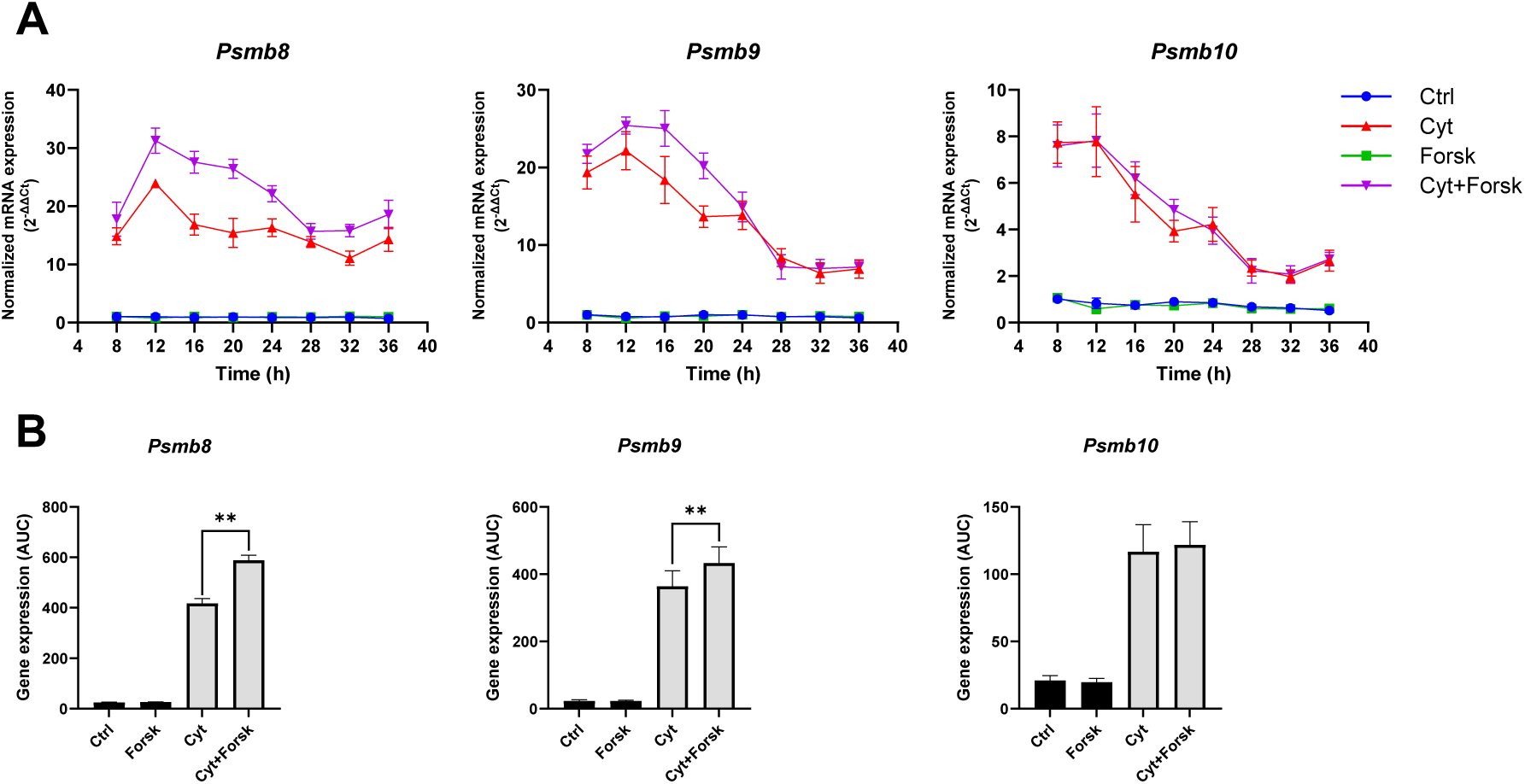
Synchronization potentiates the proinflammatory cytokine-mediated increase in ind-proteasome subunits. INS-1 cells were handled as described in FIG. 1. Normalized mRNA expression was calculated using *Hprt1* as a reference gene. **(A)** Time-dependent normalized mRNA expression of ind-proteasome subunits. **(B)** AUC of the curves in (A). Data are presented as means ± SEM (N=4). Statistics is two-tailed paired Student’s t-test.

The transcriptional activity of the BMAL1-CLOCK heterodimer and the function of the clock repressors depend on proteasomal degradation [1, 37]. Prompted by the observation that cytokine-mediated changes in clock gene mRNA expression in non-synchronized INS-1 cells were dependent on the inducible (ind)-proteasome [24], we decided to investigate the effect of synchronization on ind-proteasome subunits in the absence and presence of cytokines. By analysis of a previously published dataset [36], all three inducible subunits were found to display circadian rhythmicity in synchronized mouse islets (Table S3). The resolution of our sampling and the 8-36-hour time period studied with only one circadian cycle did not allow us to reproduce this finding by qPCR (FIG. S5). However, synchronization significantly potentiated the cytokine-mediated alterations in *Psmb8* and *Psmb9*, but not *Psmb10* expression (FIG. 3). Interestingly, bioinformatic analysis revealed that the selected core-clock proteins harbor a higher number of ind-proteasomal, compared to standard proteasomal, target sites, indicating that these proteins can be targeted by the inducible proteasomes to a higher degree than by the standard proteasome (Table S4).

In summary, *in vitro* circadian synchronization of INS1 cells potentiates the proinflammatory cytokine-mediated alterations in the expression of several core-clock mRNAs along with REV-ERBα and CLOCK proteins, and ind-proteasome subunit mRNAs.

### Cytokine-mediated core-clock gene expression is NF-κB-dependent, with circadian synchronization potentiating NF-κB signaling

Based on the observations in [24], we anticipated that the cytokine-mediated alterations in clock gene mRNA expression were NF-κB dependent. Therefore, we hypothesized that the potentiating effects of synchronization involved enhanced NF-κB signaling. To measure NF-κB activity, we investigated inducible NO synthase (*Inos/ Nos2*) expression, as its promotor contains several NF-κB *cis*-binding elements making *Inos* expression a highly sensitive NF-κB reporter. Further, NO, the product of INOS, positively feedbacks on NF-κB transcriptional activity [38]. Indeed, forskolin-induced synchronization potentiated the expression of cytokine-mediated *Inos* expression (FIG. 4A-B). As the expression of *Inos* in the investigated timeframe peaked at 8 hours, we wished to investigate an earlier time response. Synchronization potentiated *Inos* and *IκBα* expression, but not *Bmal1* at this early time-point, in cytokine-exposed INS-1 cells (FIG. S6), indicating early NF-κB activation.

**Figure 4.**
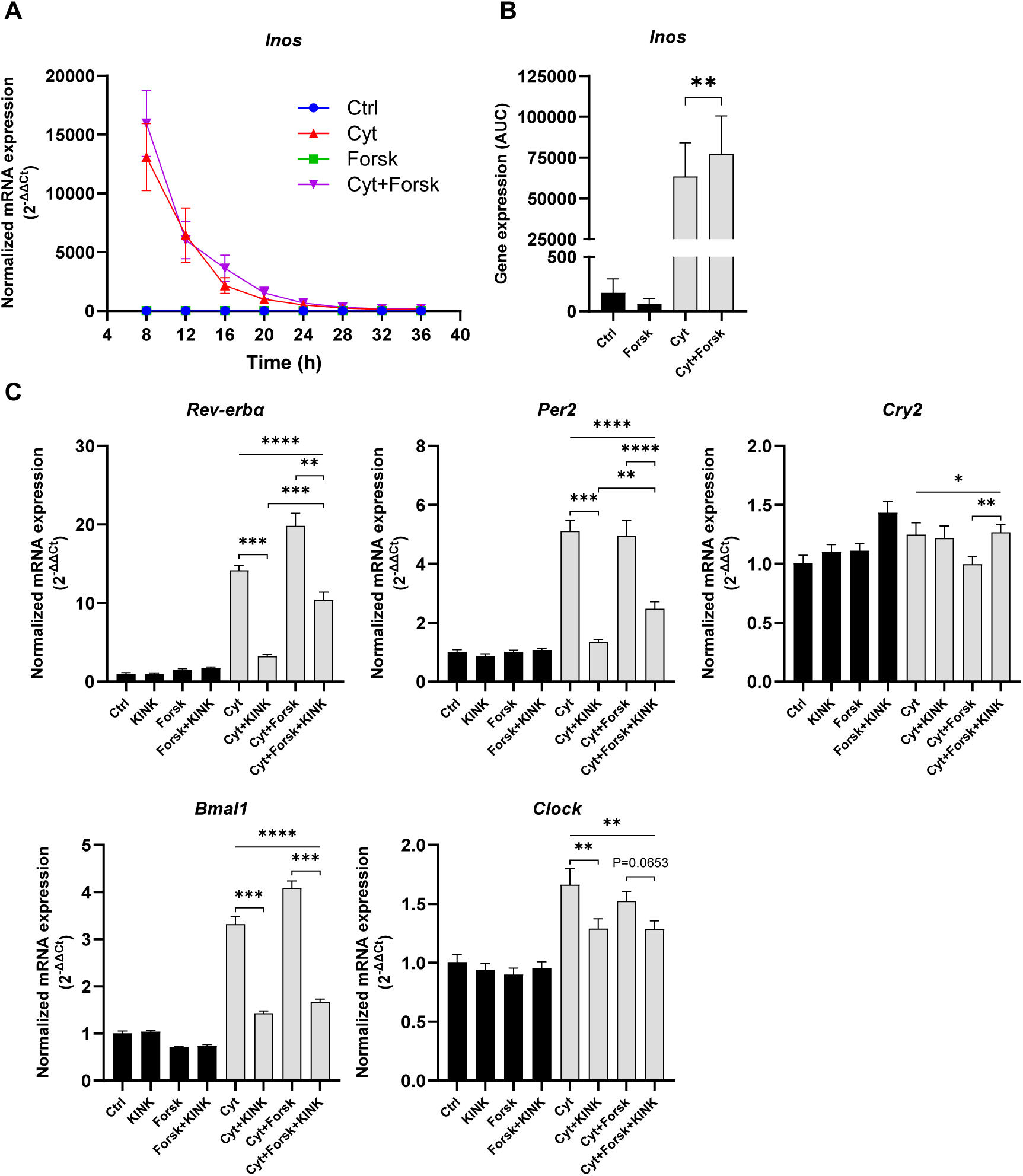
Synchronization potentiates proinflammatory cytokine-induced *Inos* expression, and NF-κB inhibition differentially reverses proinflammatory cytokine modulation of core-clock gene expression. INS-1 cells were handled as described in FIG. 1. Normalized mRNA expression was calculated using *Hprt1* and 5s rRNA as reference genes. **(A)** Time-dependent normalized mRNA expression. **(B)** AUC of the curves in (A). **(C)** Cells were additionally pre-incubated with 1 µM KINK-1 (KINK) for one hour, and exposed to the inhibitor, together with the aforementioned conditions for the 16-hour duration of the experiment. Normalized mRNA expression was calculated using *Hprt1* and 5s rRNA as reference genes. Data are presented as means ± SEM (N=4). Statistics are two-tailed paired Student’s t-test (B) and repeated measurements one-way ANOVA with p-values represented by symbols above the line and with Šídák’s corrected multiple comparisons tests (C).

Since cytokine-induced *Inos* expression was enhanced in synchronized INS-1 cells, and since iNOS inhibition abrogated cytokine-mediated alterations in clock gene mRNA expression in non-synchronized INS-cells [24], we next investigated the direct role of NF-κB in proinflammatory mediated clock gene alterations. We targeted NF-κB by a small-molecule, *IKKβ-selective Inhibitor of NF-κB* (IκB) *kinase* (IKK) inhibitor, KINK-1 [39] that by preventing phosphorylation of IκB blocks dissociation of IκB from p65 and thereby p65 nuclear translocation. One µM KINK-1, a dose selected from previous studies in our lab [32] and shown here to be non-toxic, rescued the cytokine-mediated loss of viability in non-synchronized cells as expected (FIG. S7). At 16 and 20 hours of cytokine exposure, 1 µM KINK-1 also reduced the cytokine-mediated clock gene transcript alterations approximately to baseline levels in these cells (FIG. 4C and FIG. S8). Unexpectedly, KINK-1 enhanced cytokine-induced *Psmb9* and *10* mRNA expression in non-synchronized cells, suggesting the existence of NF-κB dependent repressors of the expression of ind-proteasomal components (FIG. S9A).

In synchronized cells, NF-κB inhibition differentially attenuated cytokine modulation of core-clock transcripts (FIG. 4C and FIG. S8) and ind-proteasomal subunit expression (FIG. S9A), with the effect of KINK-1 in cytokine-exposed cells differing depending on the transcript in question. For *Rev-erbα*, *Per2*, and *Bmal1*, NF-κB inhibition reversed cytokine-mediated alterations at both 16 and 20 hours, with the same trend (p=0,065) for *Clock* at 16 hours (FIG. 4C and FIG. S8). Synchronization reduced *Cry2* expression relative to the other conditions, and this inhibition was reversed by KINK-1 at 16 hours.

KINK-1 exposure further increased cytokine-induced expression of *Psmb10* at both 16 and 20 hours, and of *Psmb9* at 20 hours with statistical trends at 16 hours in both non-synchronized and synchronized cells (FIG. S9A). *Psmb8* followed the pattern for *Psmb10* and -*9* with regard to the potentiating effect of KINK-1 on cytokine-induced expression in non-synchronized cells, but with the opposite pattern in synchronized cells as indicated by the overall significant ANOVA. However, these differences were not significant in post-hoc t-tests.

To better appreciate the impact of KINK-1 on cytokine-mediated alterations of clock and proteasome mRNAs in synchronized versus non-synchronized cells, the percentage change in expression caused by NF-κB inhibition during cytokine exposure was calculated, correcting for the baseline expression in non-cytokine exposed cells (FIG. S9B). In general, the relative effect sizes of NF-κB inhibition on the expression of both clock and ind-proteasomal genes were attenuated in synchronized versus non-synchronized cells (FIG. S9B).

To summarize, NF-κB inhibition differentially attenuates cytokine-mediated alterations in core-clock gene transcripts and ind-proteasomal subunit expression.

### Synchronization potentiates cytokine-induced β-cell endoplasmic reticulum stress and the intrinsic apoptotic pathway

NO is a key conveyor of cytokine-mediated ER stress, since NO inhibits the Sarcoplasmic/endoplasmic reticulum Ca^2+^-ATPase (SERCA) pump, causing ER Ca^2+^ depletion [20]. Therefore, we explored if ER stress markers were affected by synchronization of cytokine-exposed INS-1 cells. Indeed, cytokine-induced *Xbp1* splicing *(sXbp1*), as well as total and unspliced *Xbp1*, *Bip,* and *Atf4* expression (FIG. 5 and FIG. S10) were all potentiated by synchronization.

**Figure 5.**
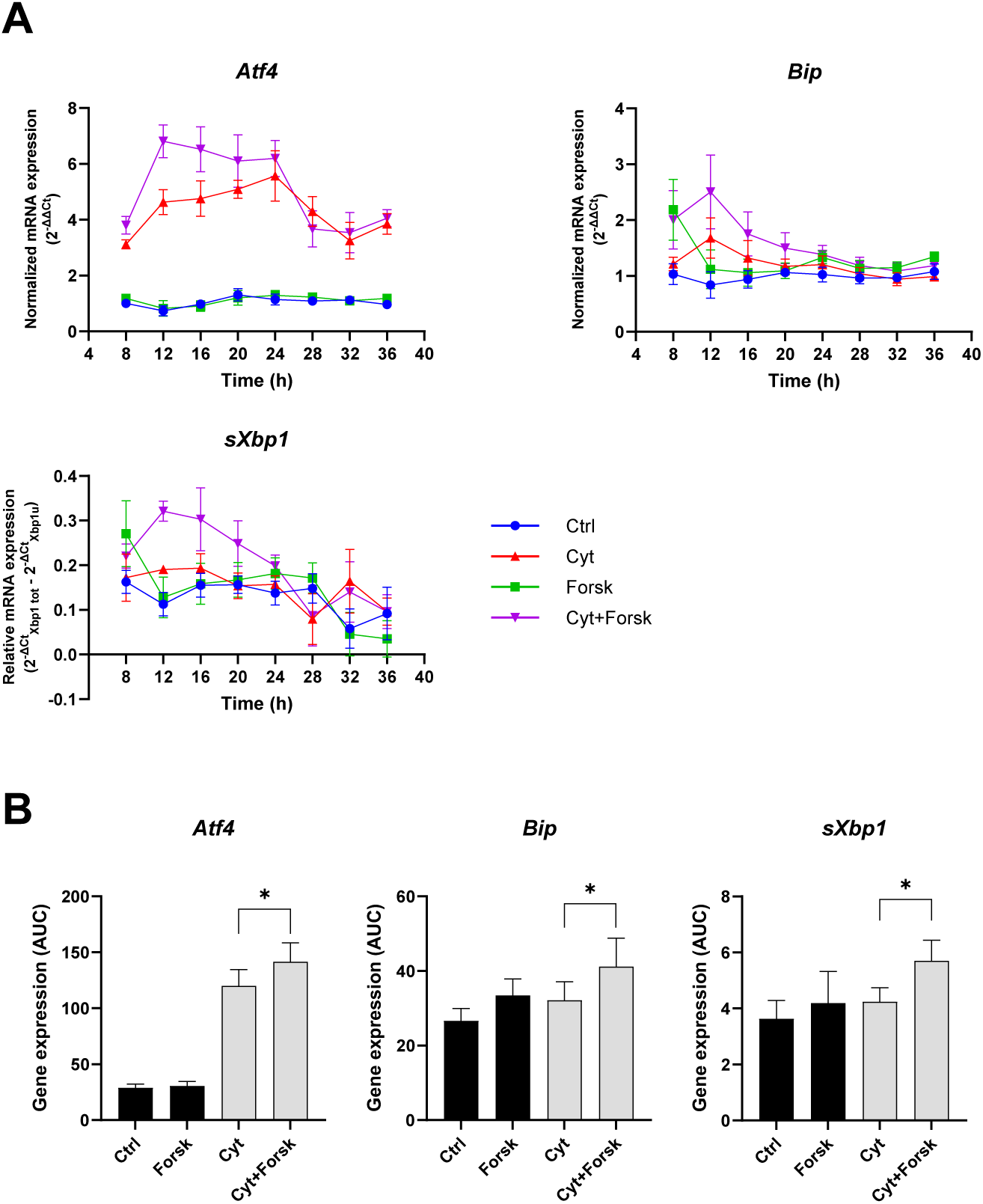
Synchronization potentiates the proinflammatory cytokine-mediated increases in ER stress-associated genes. INS-1 cells were handled as described in FIG. 1. Normalized mRNA expression was calculated using *Hprt1* and 5s rRNA as reference genes. **(A)** Time-dependent normalized mRNA expression. **(B)** AUC of the curves in (A). Data are presented as means ± SEM (N=4). Statistics is two-tailed paired Student’s t-test.

The cytokine-induced ER stress response is believed to contribute to activation of the intrinsic (mitochondrial), but not by the extrinsic (death domain) apoptotic pathway [18]. Cytokine exposure of synchronized cells was associated with increased activity of intrinsically activated Caspase 9 but not extrinsically activated Caspase 8 activity (FIG. S11A), resulting in a small but significant increase in cytotoxicity (apoptosis and late apoptotic necrosis) (FIG. S11B+C).

In summary, synchronization potentiates the cytokine-mediated activation of ER stress and the intrinsic apoptotic pathway.

### NF-κB inhibition differentially affects cytokine toxicity in a cytokine concentration-dependent manner

Having demonstrated the importance of NF-κB signaling in cytokine-mediated clock gene transcript and ind-proteasomal subunit expressions as well as in the effects of synchronization on β-cell viability, we next tested the role of NF-κB in cytokine-mediated cytotoxicity in insulin-producing cells.

In the absence of cytokines, synchronized cells were more viable than their non-synchronized counterpart, and KINK-1 attenuated this advantage, albeit not completely (FIG. S12). We next verified that NF-κB inhibition reduced cytokine-mediated cytotoxicity in the INS-1 Per2-Luc reporter cells exposed to varying IL-1β concentrations with a fixed IFN-γ concentration.

Indeed, KINK-1 reduced the IL-1β dose- and time-dependent increase in apoptosis and post-apoptotic necrosis in both non-synchronized and synchronized cells (FIG. 6A+B and S12). At 10 pg/mL IL-1β KINK-1 increased cytotoxicity in non-synchronized cells suggesting a protective role of NF-κB at low IL-1β concentrations in these cells, while KINK-1 reduced cytotoxicity in synchronized cells (FIG. S12).

**Figure 6.**
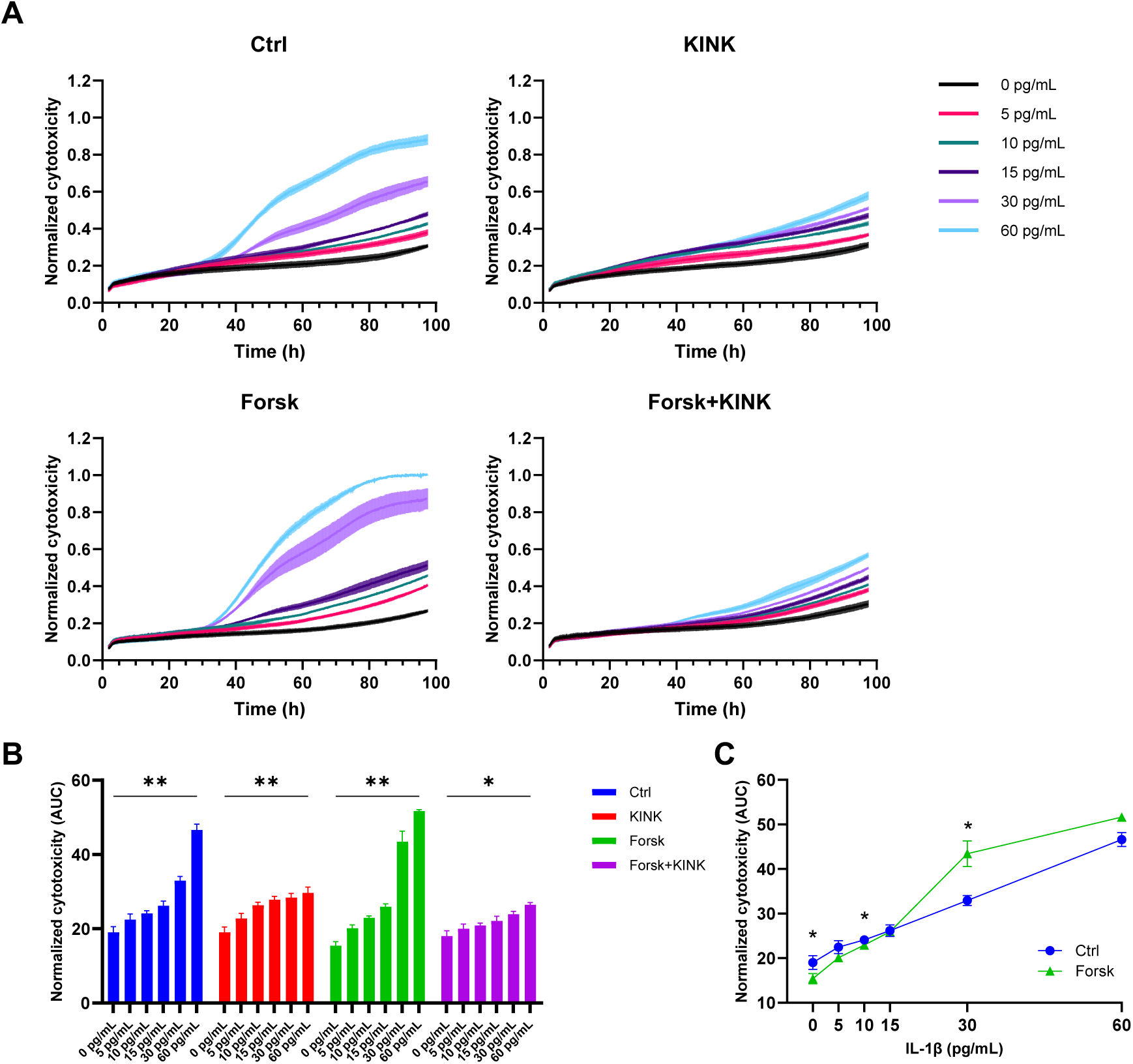
NF-κB inhibition reduces the dose-dependent proinflammatory cytokine-mediated cytotoxicity in both synchronized and non-synchronized cells, while synchronization dose-dependently modifies cytokine-mediated cytotoxicity. INS-1 Per2-Luc cells were cultured in the presence of varying concentrations of mIL-1β (legend) and 0.1 ng/ml rIFN-γ. Cells were synchronized with a one-hour 10 μM forskolin pulse (Forsk), as well as 1 µM KINK-1 (KINK) in a one-hour preincubation and for the duration of the experiment. Cells not exposed to forskolin or KINK-1 have been annotated as control (Ctrl). **(A)** Cytotoxic response normalized to the overall maximum signal obtained. Cytotoxicity was monitored in real-time at 10-minute intervals for 96 hours. **(B)** AUC of the normalized cytotoxicity response. **(C)** Dose-response of cytokine concentrations on AUC of normalized cytotoxicity response. Data are presented as means ± SEM (N=3). Statistics are repeated measurements one-way ANOVA with p-values represented by symbols above the line (B) and two-tailed paired Student’s t-test (C).

While the effects of synchronization and NF-kB inhibition were neutral at 15 pg/ml of IL-1β, synchronization potentiated cytokine-induced apoptosis and late apoptotic necrosis at 30 and 60 pg/mL IL-1β (FIG. 6 and S12). When these data were plotted as a cytokine dose-response rather than a time-response relationship, the response was bimodal, since synchronization reduced cytotoxicity at low concentrations of IL-1β (5-10 pg/mL), while enhancing toxicity at higher concentrations (≥30 pg/mL), with the apparent turning point being at 15 pg/mL (FIG. 6C).

We intended to verify key findings from the above experiments in the human EndoC-βH3 cell line [33]. However, even at much higher cytokine concentrations, these cells were not cytokine-sensitive (FIG. S13).

### NF-κB inhibition differentially affects cytokine-mediated changes in period and acrophase in a cytokine concentration-dependent manner

We next wished to investigate the dose-dependent effect of cytokines and NF-κB inhibition on circadian parameters using the INS-1 Per2-Luc reporter cells (FIG. 7 and FIG. S14+S15). Non-detrended traces demonstrated that high concentrations of IL-1β (30 and 60 pg/mL) reduced Per2-Luc bioluminescence at 96 hours, with the decrease starting after approx. 10 hours, which was reversed by KINK-1 (FIG. S14). This reduction was associated with a decrease in *Per2* promotor-driven *Luc* mRNA at 20 hours, but not in *Per2* mRNA (FIG S16), suggesting that the decreased bioluminescence signal is indeed due to reduced expression of luciferase as an expression of general effects on circadian rhythmicity, concurrent with the increased *Per2* expression which would provide a repressive function on the core-clock. Hence, the reduced bioluminescence signal is not an artifact caused by a global effect of cytokine toxicity, but an expression of circadian amplitude. This notion was supported by the intact oscillations (FIG. 7 and S14) and the toxicity data (FIG. 6).

**Figure 7.**
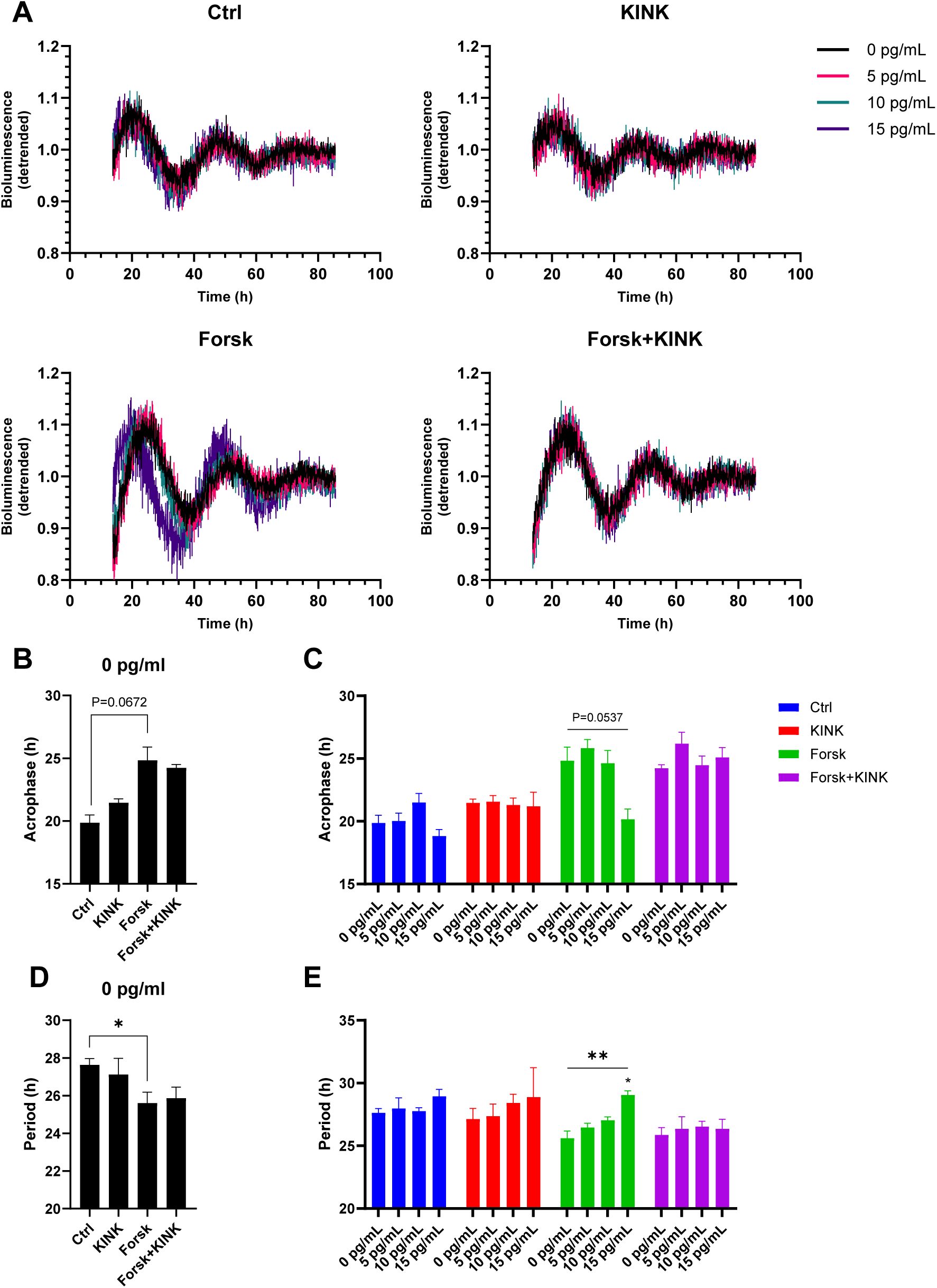
NF-κB inhibition normalizes the dose-dependent proinflammatory cytokine-mediated changes in acrophase and period. INS-1 Per2-luc cells were handled as described in FIG. 6 **(A)** Detrended Per2-Luc bioluminescence signal, recorded at 10-min intervals presented in the functional window (13.5-85.5 hours) of the analysis, from which acrophase in the absence **(B)** and presence **(C)** of cytokines, in addition to period in the absence **(D)** and presence **(E)** of cytokines, have been determined. Data are presented as means ± SEM (N=3). Statistics are two-tailed paired Student’s t-test for (B) and (D), and repeated measurements one-way ANOVA with p-values represented by symbols above the line and with Dunnett’s multiple comparisons test to 0 pg/mL for (C) and (E).

Non-synchronized cells displayed inherent rhythmicity (FIG. 7A) from which acrophase (∼20 hours) (FIG. 7B) and period (∼27 hours) (FIG. 7D) could be determined after detrending. Indeed, entrainment by forskolin markedly altered the rhythmicity of INS-1 cells closer to expected circadian period (FIG. 7B+D) (acrophase and period both ∼25 hours). Interestingly, cytokines dose-dependently shortened the acrophase to ∼20 hours and prolonged the period to ∼30 h in synchronized but not in non-synchronized cells (FIG. 7C+E). Notably, NF-κB inhibition normalized the cytokine-modulation of acrophase and period in synchronized INS-1 Per2-Luc cells at low (≤15 pg/mL) IL-1β concentrations.

At 30 pg/mL of IL-1β, KINK-1 failed to reverse the cytokine-mediated period lengthening, yet synchronized cells maintained a rhythmic profile of the detrended traces (FIG. 8). At 60 pg/mL, no further period lengthening was observed, coinciding with plateauing of the maximal cytotoxic response (FIG. 6A). For synchronized cells exposed to 60 pg/mL IL-1β and KINK-1, the period length reached the upper limit (34 hours) of the period length window used in the analysis (18-34 hours) (FIG. 8D, dotted line), with a flat profile of the detrended trace (FIG. 8C), indicating that rhythmicity in synchronized cells treated with KINK-1 in the presence of 60 pg/ml of IL-1β was lost or had become infradian. Interestingly, these cells still suffered less apoptosis and post-apoptotic necrosis than cells not treated with the NF-κB inhibitor (FIG. 6A+B), suggesting a protective effect of NF-κB inhibition against cytokine toxicity uncoupled from clock rescue.

**Figure 8.**
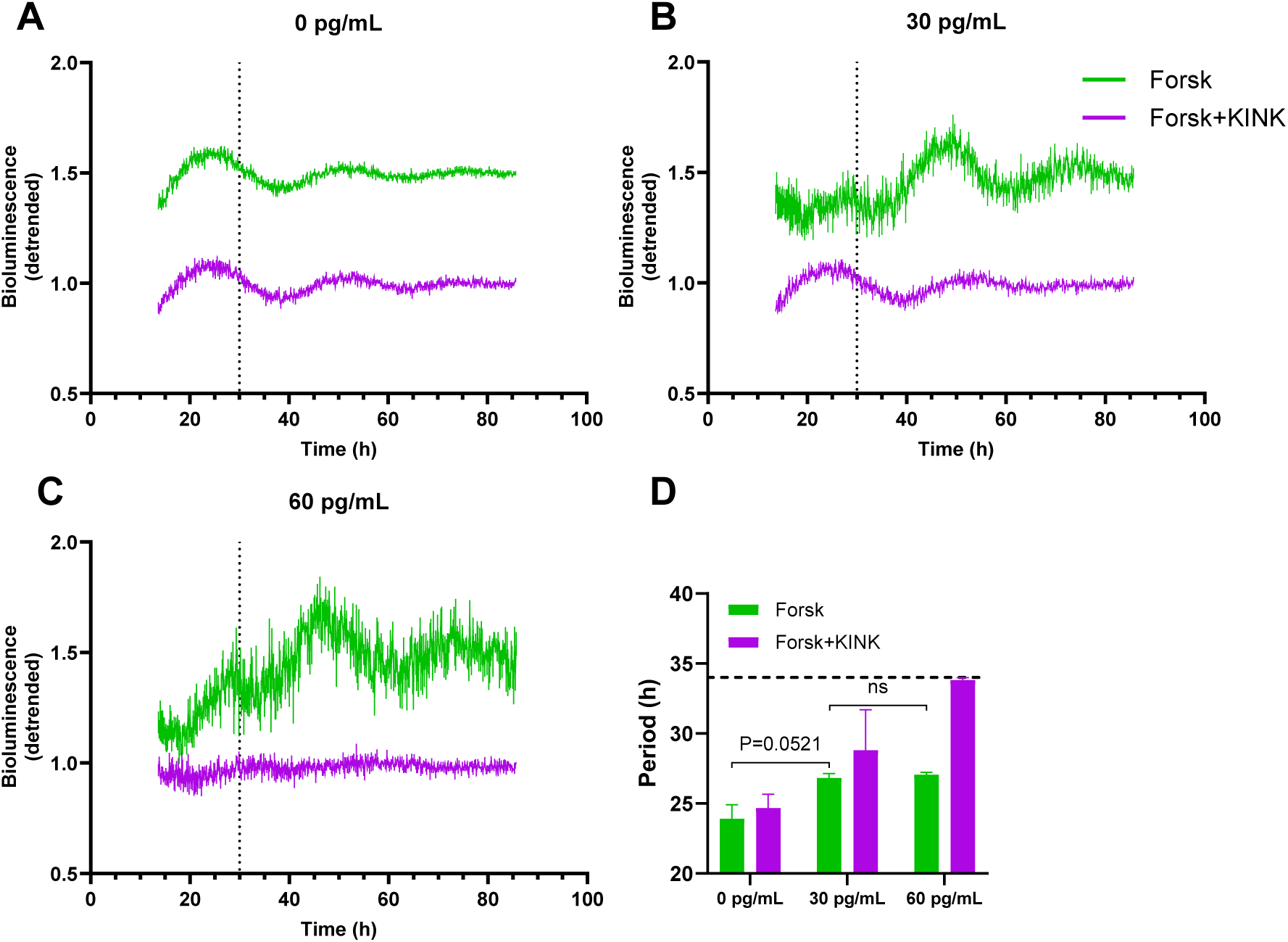
NF-κB inhibition fails to normalize period length at cytotoxic IL-1β concentrations. INS-1 Per2-luc cells were synchronized with a one-hour 10 μM forskolin pulse (Forsk) and 1 µM KINK-1 (KINK) in a one-hour preincubation and for the duration of the experiment. Cells were cultured in the absence **(A)** presence of 30 **(B)** or 60 **(C)** pg/mL mIL-1β with 0.1 ng/ml rIFN-γ, with bioluminescence recorded at 10-min intervals and data detrended. **(D)** Period length of (A-C). Period length was determined using detrended data in an analysis window between 30 and 85 hours (vertical dotted line, A-C), with an estimated period length between 18 and 34 hours (horizontal dotted line, D). Data are presented as means ± SEM (N=3). Statistics is two-tailed paired Student’s t-test.

In summary, IL-1β and IFN-γ dose-dependently advance acrophase and lengthen the period in the non-cytotoxic cytokine concentration range in an NF-κB dependent manner. In contrast, at cytotoxic cytokine concentrations, the increased period length plateaus, with NF-κB inhibition being unable to reverse this effect and causing loss of circadian rhythmicity.

## Discussion

Our finding that a combination of proinflammatory cytokines, IL-1β and IFN-γ, alters the expression level of several core-clock components at the protein level expands our earlier reporting of cytokine-induced changes in the transcription of these components [24]. This observation is important for a comprehensive interpretation of how cytokines regulate the transcriptional-translational feedback loop in the clock machinery, as clock proteins are the transcriptional repressors and activators. Since the previous intriguing observations were obtained in a non-synchronized cell system, we here investigated cytokine regulation of clock components at both the transcriptional and translational level in a forskolin-synchronized cell model.

We demonstrate that circadian synchronization significantly potentiated cytokine-mediated alterations of the transcription of several core-clock genes and, interestingly, the level of a more restricted repertoire of clock proteins (the activator CLOCK and the repressor PER2). Since circadian dynamics depend on proteasomal degradation of clock proteins, we anticipated parallel regulation of proteasomal components. Synchronization did indeed potentiate the cytokine-induced transcription of the ind-proteasomal subunits *Psmb8* and *Psmb9*.

Further mechanistic studies revealed that synchronization potentiated the cytotoxic effect of proinflammatory cytokines in a cytokine dose-dependent manner, associated with potentiation of ER stress markers, *Inos* mRNA expression, and increased Caspase 9 activity. We reasoned that cytokine-triggered NF-κB signaling might be a unifying signaling pathway conveying synchronization-dependent potentiation of cytokine-regulation of clock and ind-proteasomal component expression, as well as the summarized mechanistic pathways. Indeed, pharmacological inhibition of NF-κB activity reverted the cytokine-mediated alterations in clock gene expression in both non-synchronized and, to a lesser extent, synchronized cells, associated with reduced cytotoxicity. In accordance with the known bimodal effect of proinflammatory cytokines on β-cell function and viability [40, 41], NF-κB inhibition normalized acrophase advancement and period lengthening caused by low (≤15 pg/mL) IL-1β. At high (≥30 pg/mL) IL-1β, NF-κB inhibition failed to revert period lengthening and results in loss of circadian rhythmicity.

Taken together, our data suggest that proinflammatory cytokines perturb the circadian clock in an NF-κB dependent manner and that clock perturbation desensitizes insulin-producing cells to cytokine-mediated assault, likely as part of an integrated anti-inflammatory β-cell defense response.

The observation that NF-κB is crucial for both the cytokine-mediated uncoordinated change in clock gene expression and in perturbing circadian rhythmicity supports our earlier findings that cytokine-activation of multiple pathways down-stream of NF-κB is associated with the cytokine-mediated change in core-clock gene expression profile [24]. The involvement of NF-κB signaling in core-clock gene expression is supported by the observation that multiple clock genes have NF-κB binding sites. Importantly, p65 itself displays an oscillatory protein expression and activity profile, required for circadian oscillations of core-clock genes and proteins in non-cytokine exposed mouse embryonic fibroblasts (MEF), and a functional IKK complex is required for mouse locomotor rhythms [22, 23, 42]. In synchronized human U2OS cells, upregulation of p65 shortened period length, while inhibition increased period length, indicative of an inhibitory function by p65 on BMAL1-CLOCK transcriptional activity, similar to the repressor function of CRY1 [23]. This contrasts with our findings that NF-κB inhibition in non-cytokine-exposed synchronized INS-1 cells did not change rhythmic parameters, and inhibition of NF-κB in cytokine-exposed synchronized cells shortened the period. The discrepancy could be an indication of cell-specific differences in NF-κB action as reflected, for example, by the anti-apoptotic action of NF-κB in the liver vs the pro-apoptotic effects in the β-cell [43, 44]. A cell-specific NF-κB action is further illustrated in murine liver, where lipopolysaccharide (LPS) exposure increases p65 binding and activity resulting in inhibition of clock repressors, but not activators [22], in contrast to the general NF-κB dependent upregulation observed in our study. Additionally, LPS exposure results in genome-wide relocalization of BMAL1-CLOCK to sites in proximity to genes involved in inflammatory and metabolic signaling and apoptosis and is dependent on functional p65 [22]. This concept may explain the NF-κB-dependent cytokine-mediated effects on BMAL1-CLOCK-controlled genes shown here.

Interestingly, the rescue potential of NF-κB inhibition on rhythmic parameters depended on the IL-1β concentration. At non-toxic cytokine concentrations, incapable of altering clock gene expression in non-synchronized cells (≤15 pg/mL IL-1β) [24], NF-κB inhibition reversed cytokine-mediated alterations in rhythmicity (FIG. 8 and 9). In contrast, at higher cytotoxic levels of IL-1β (≥30 pg/mL), NF-κB inhibition restrained the reintroduction of a rhythmic behavior, while still reducing cytokine-mediated cytotoxicity (FIG. 8 and 10). The uncoupling of the effects of the KINK-1 inhibitor on clock perturbation and toxicity may be caused by the global NF-κB inhibiting activity of KINK-1, unselective for NK-κB homo- or heterodimers, the balance of which may finetune the cellular responses to inflammatory stimuli.

Synchronized cells exposed to high cytokine concentrations in the absence of the NF-κB inhibitor were able to reintroduce rhythmicity after the first ∼30 hours of non-rhythmic behavior, coinciding with the uncoordinated clock transcriptional changes (FIG. 2) reversible by NF-κB inhibition (FIG. 4 and S8), possibly as a defensive response.

The potential involvement of the ind-proteasome in regulation of the circadian transcriptional-translational feedback-loop is supported by our earlier observation that inhibiting the ind-proteasome reverses the cytokine-mediated effects on clock gene expression [24]. Taken together with the observation that the ind-proteasome is constitutively expressed and active in β-cells and that the activity of the ind-proteasome is induced by IFN-γ or low concentrations of IL-1β [45, 46], our data suggests that the ind-proteasome regulates the dynamics of the molecular β-cell clock, an action potentiated by synchronization. This notion is exemplified by the discrepancy between *Bmal1* mRNA and protein (FIG. 1 and 2)

REV-ERBα mRNA and protein expressions were cytokine-coregulated and both potentiated by synchronization. Pharmacological activation of REV-ERBα impairs autophagy and insulin expression and secretion while promoting apoptosis in β-cells, likely through clock-independent transcriptional effects [24, 47]. The loss of β-cell function and increased apoptosis might be an effect of impaired autophagy, as loss of autophagy results in loss of β-cell mass in mice [48]. Accordingly, REV-ERBα/β agonists impair autophagy and induce apoptosis in a range of cancer cells [49], supporting REV-ERBα/β’s role in apoptosis. Thus, the potentiated cytokine-mediated cytotoxicity during synchronization could in part be explained by the potentiated expression levels of REV-ERBα in synchronized cells. However, it cannot be rejected that these REV-ERBα mediated effects are clock-independent.

The widely-used one-hour forskolin-pulse was used here as a synchronization stimulus, shown to upregulate *Per1/2* [24] which is known to result in entrainment of the molecular clock [28]. Forskolin elevates cAMP, which increases Protein kinase A (PKA) activity, resulting in Ca^2+^/cAMP responsive element binding protein (CREB) activity, and thereby *Per1/2* expression [28], but also of numerous other genes. We do not believe that our observations are confounded by off-clock target effects, for the following reasons. First, cAMP has a half-life of <1 ½ minutes [50], resulting in a very transient increase in cAMP after a one-hour forskolin pulse. This is supported by the observation that insulin secretion from murine islets exposed to 10 µM IBMX and 1 µM forskolin increased ∼350 % during a 40 min exposure, which normalized 30 min after the cessation of IBMX and forskolin exposure [51], indicating cAMP turnover in β-cells is very fast to enable minute-by-minute adjustments in insulin secretion. As cytokine-mediated β-cytotoxicity is observed after 24-96 hours, it is unlikely that these effects are due to constant high cAMP levels. In fact, cAMP protects β-cells against apoptosis [52], similar to what is seen in this current study in the absence of proinflammatory stress. Second, cAMP can both inhibit and promote NF-κB signaling, depending on stimulus and cell type [53]. However, in non-cytokine exposed cells, NF-κB signaling in INS-1 cells was unchanged at one [24] and three hours (FIG. S8) post-forskolin pulsing, supporting that the potentiating effects of synchronization on cytokine-action are not due to off-target effects secondary to transiently increased cAMP.

Non-forskolin pulsed cells displayed a weak rhythmic profile including an early Per2-Luc spike (FIG. 7 and S2) likely due to the experimental handling including temperature shifts, plating, and medium change providing a minor synchronization signal [54]. However, forskolin exposure caused a change in both acrophase and period length direction of the expected parameters compared to non-forskolin exposed cells, indicating that the effects of cytokines depend on the robustness of the synchronization.

A major strength of the methodological approach in this study was the use of rhythmicity reporter cells multiplexed to a cytotoxicity assay, as this gives parallel insight into the association between rhythmicity and cytotoxicity in the same cell population. In combination with extensive qPCR and Western blotting verification of fluctuations in core-clock components, this assay enabled interpretation of descriptive and mechanistic data.

A weakness is the use of INS-1 cells as the only model system, which is not just related to the inherent limitations of immortalized rodent cell lines, but also related to the limitation of using rapidly dividing cells for circadian monitoring. Even after entrainment, INS-1 cell oscillations were dampened after approximately three cycles, limiting the number of cycles available to calculate rhythmic parameters. This is especially the case for INS-1 Per2-luc cells exposed to high levels of IL-1β, where the first 30 hours could not be used for analysis, limiting the oscillations used in the analysis to two cycles.

The potential use of the human β-cell line EndoC-βH3 was explored in this study but had to be abandoned since these cells were found to be unresponsive to proinflammatory cytokines, a problem also encountered by other laboratories (Soleimanpour S, personal communication). Thus, our observations should be repeated in primary islets, preferably human.

Having observed that desynchronization desensitizes insulin-producing cells to inflammatory assault and protects against selected cytokine-mediated gene expression alterations, investigation of global cytokine-mediated transcriptomic and proteomic effects in synchronized vs non-synchronized β-cells is warranted. We speculate that the proinflammatory transcriptional activity of NF-κB in synchronized cells is enhanced by interaction with clock transcription factors, as it has been suggested for the clock activator *BMAL1* [22], and since desynchronization provides a defensive response, similar to what has been shown for β-cell phenotypic de-differentiation in response to cytokine assault [55, 56]. This notion should also be explored in the context of other β-cell stressors such as glucolipotoxicity and in other cell-types. Finally, the bimodal action of NF-κB depending on proinflammatory stress intensity and the disparate effects on clock rhythmicity and cell-toxicity require more investigation.

In conclusion, our data suggest that proinflammatory cytokines perturb the circadian clock in an NF-κB dependent manner and that clock perturbation desensitizes insulin–producing cells to cytokine-mediated assault, likely as part of an integrated anti-inflammatory β-cell antiapoptotic defense response.

## Acknowledgments

We thank Frederik Vilhardt and Jonas Hyld Steffen (University of Copenhagen, Copenhagen, Denmark) for their great help in establishment of the INS-1 *Per2-Luc* cell line. We further thank Charna Dibner and Volodymyr Petrenko (University of Geneva, Geneva, Switzerland) for their expert technical assistance and helpful discussions.

## Abbreviations

Bmal1: Brain and muscle arnt-like 1
Clock: Circadian locomotor output cycles kaput
CREB: Ca^2+^/cAMP responsive element binding protein
Cry: Cryptochrome
ER: endoplasmic reticulum
HDAC3: Histone deacetylase 3
IFN-γ: Interferon-γ
IKK: IκB kinase
IL-1β: Interleukin-1β
iNOS: Inducible nitric oxide synthase
IκB: Inhibitor of NF-κB
LPS: lipopolysaccharide
MAPK: Mitogen activated protein kinase
NF-κB: Nuclear factor of kappa light polypeptide gene enhancer in B-cells
Per: Period
Per2-Luc: Period2 promotor-driven Luciferase
PKA: Protein kinase A
Rev-erbα or Nr1d1: Nuclear receptor subfamily 1 group D member 1
SCN: Suprachiasmatic nucleus
SERCA: Sarcoplasmic/endoplasmic reticulum Ca^2+^-ATPase

## Supplementary figures

**Figure S1.**
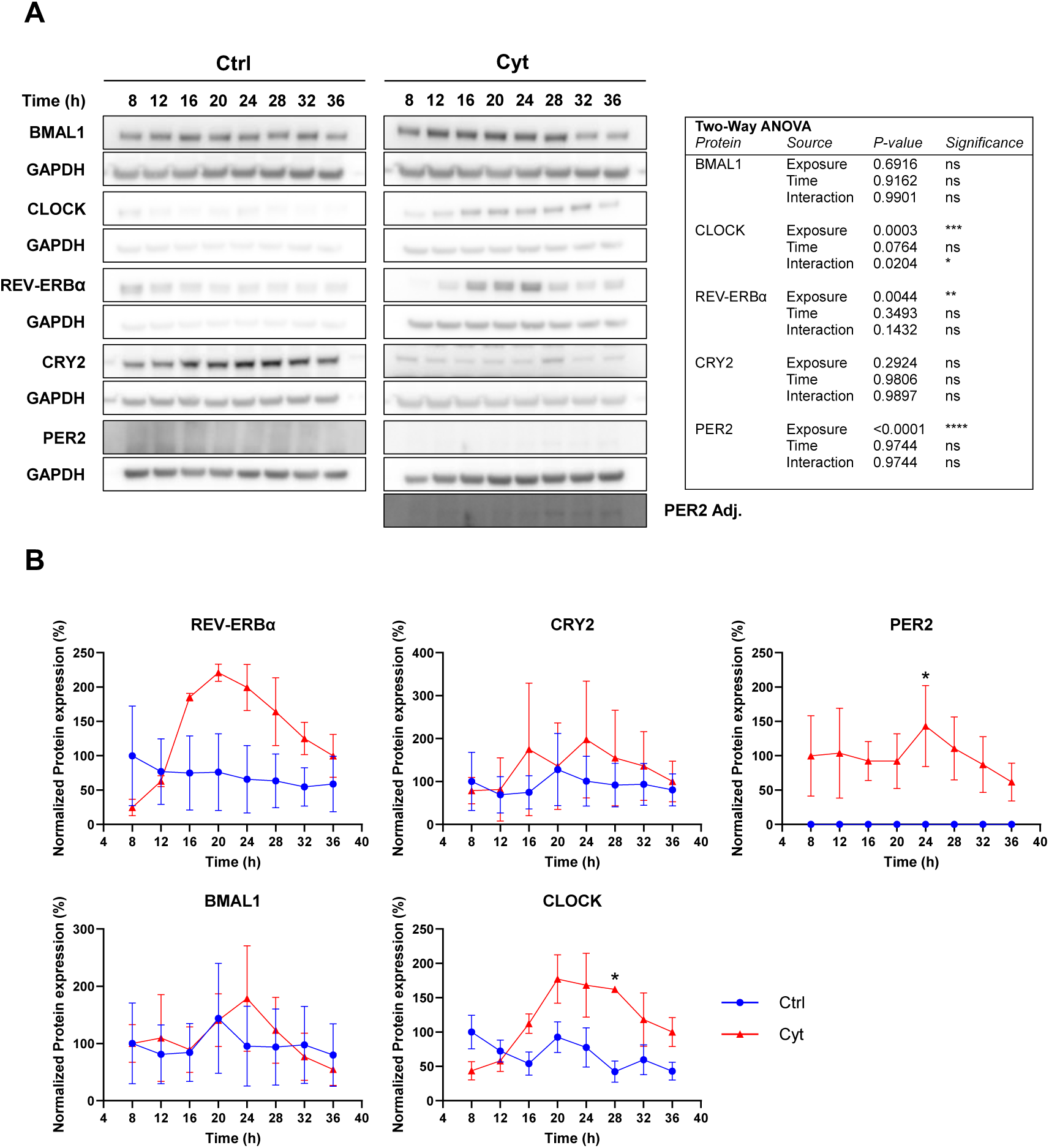
Proinflammatory cytokines induce core-clock protein expression. INS-1 cells were exposed to culture medium (Ctrl) or cytokines (Cyt) described in FIG. 1. **(A)** Samples were analyzed by SDS-PAGE and Western blotting with GAPDH as the internal loading control. PER2 Adj. is the PER2 band gamma adjusted to allow for illustration of the bands, while the unadjusted PER2 bands have been used for quantification **(B)** Quantification and normalization of the protein expression levels. Data are presented as means ± SEM (N=2-4). Statistics is two-way ANOVA with Šídák’s corrected multiple comparisons tests. A summary of the statistical tests is presented in the table.

**Figure S2.**
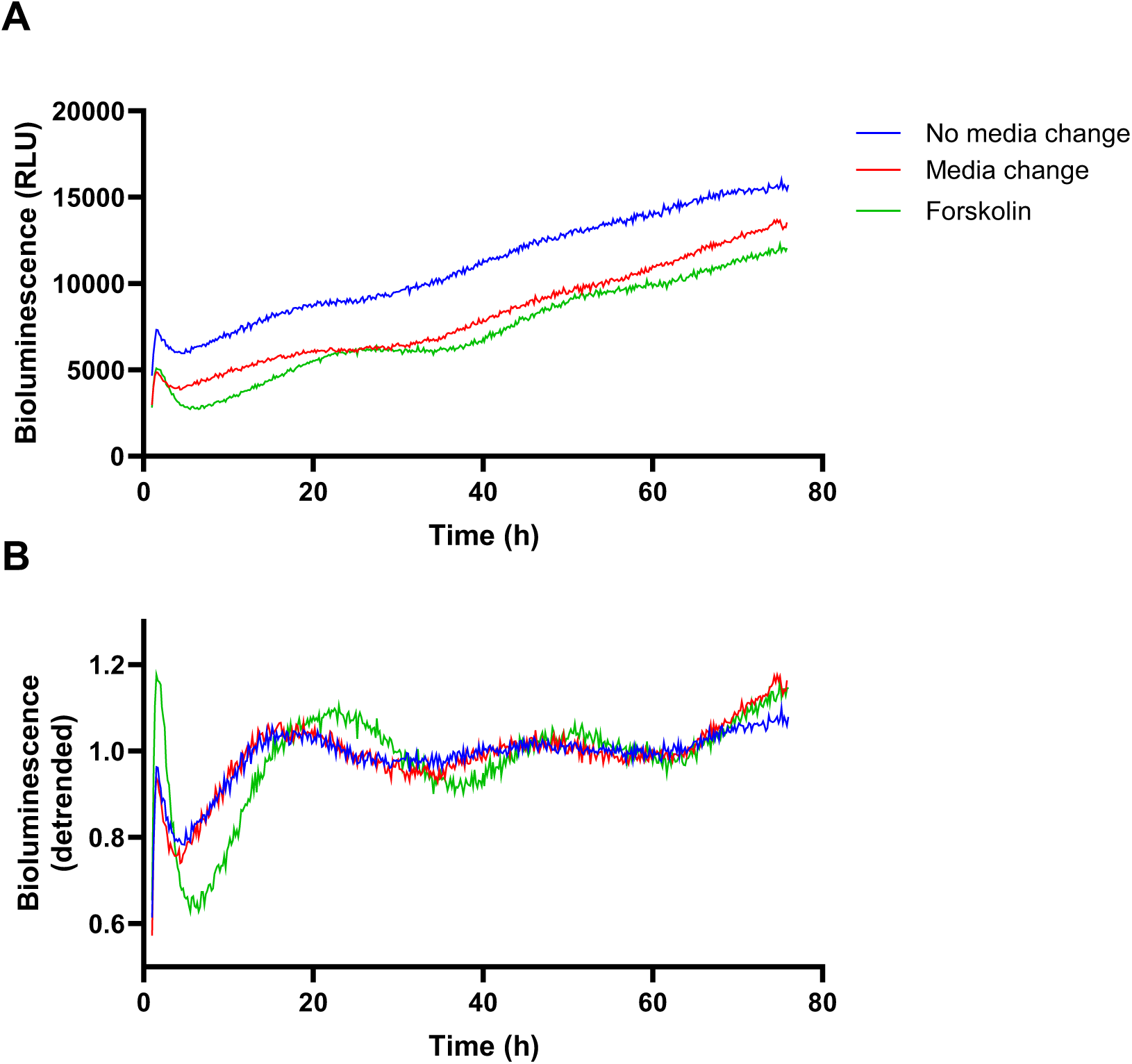
Forskolin induces potent INS-1 Per2-Luc oscillations. INS-1 Per2-Luc cells were plated and incubated for 3 days before exposure one hour 10 µM forskolin pulse (Forskolin). Additionally, cells were subjected to experimental medium change (medium change), or direct addition of luciferin (no medium change). After forskolin stimulation, 0.1 mM D-luciferin was added, and bioluminescence was monitored at 10-min intervals for 72 hours **(A)**, with detrended values using a moving 24-hour average in **(B)**. Data are presented as means of 3 technical replicates (N=1).

**Figure S3.**
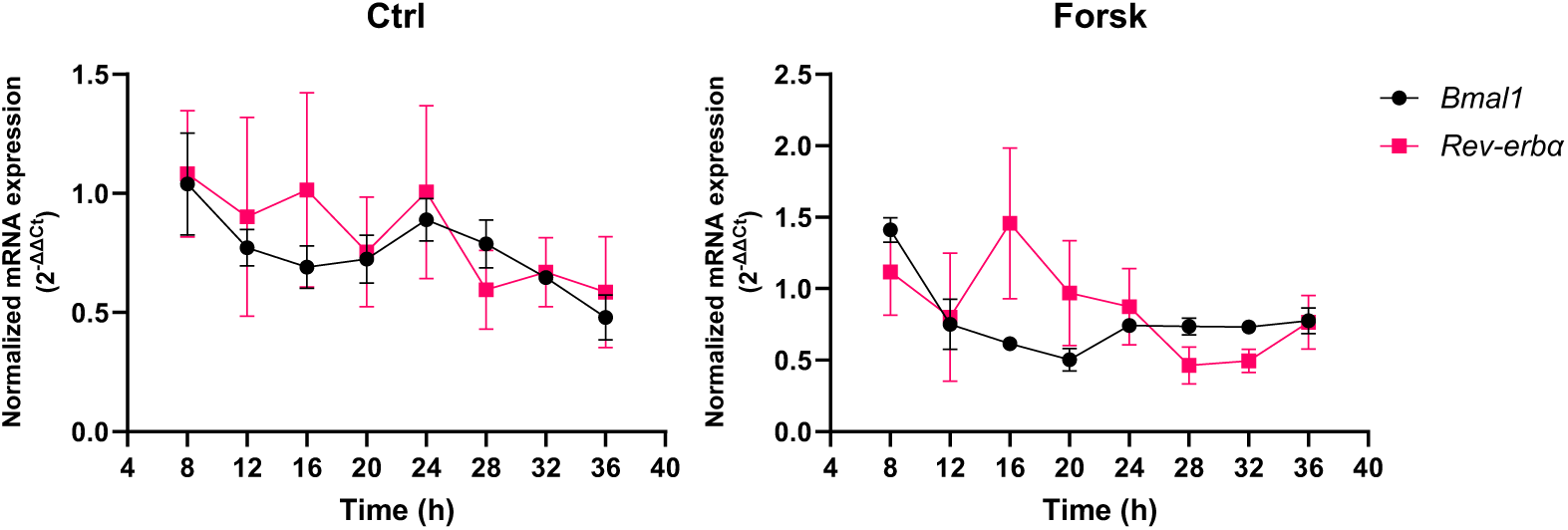
Forskolin pulsing results in antiphasic expression of core-clock genes. INS-1 cells were exposed to a one hour 10 µM forskolin pulse (Forsk), or no pulse (Ctrl). Cells were harvested at 4-hour intervals for between 8 and 36 hours. Relative mRNA expression is calculated using *Hprt1* and 5s rRNA as reference genes. Data are presented as means ± SEM (N=4).

**Figure S4.**
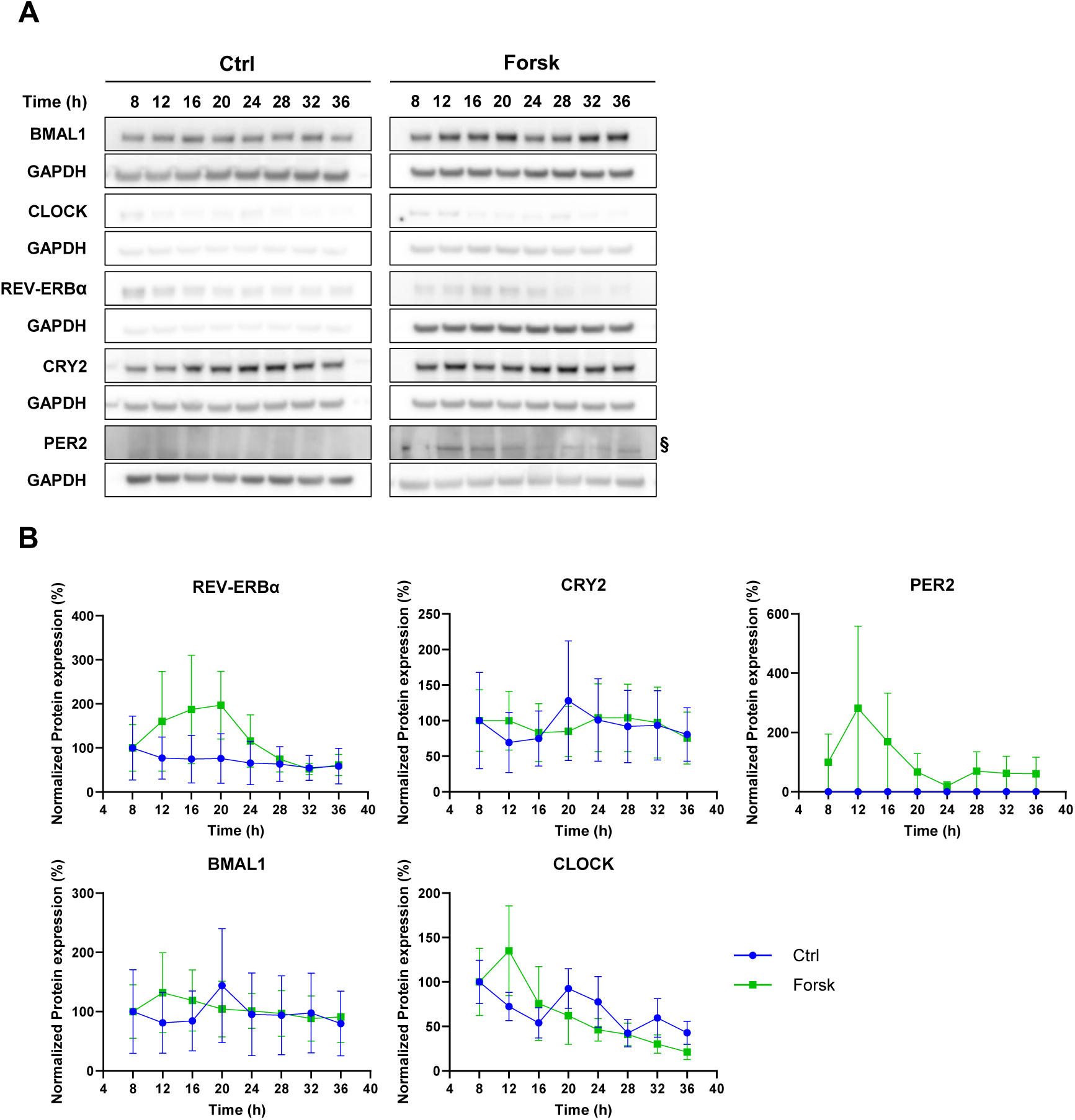
Forskolin pulsing results in rhythmic expression of core-clock proteins. INS-1 cells were handled as described in FIG. S3. **(A)** Samples were analyzed by SDS-PAGE and Western blotting with GAPDH as the internal loading control. **(B)** Quantification and normalization of the protein expression levels. Data are presented as means ± SEM (N=2-4). §: only one of four blots showed detectable PER2 bands, illustrated by the blot in A.

**Figure S5.**
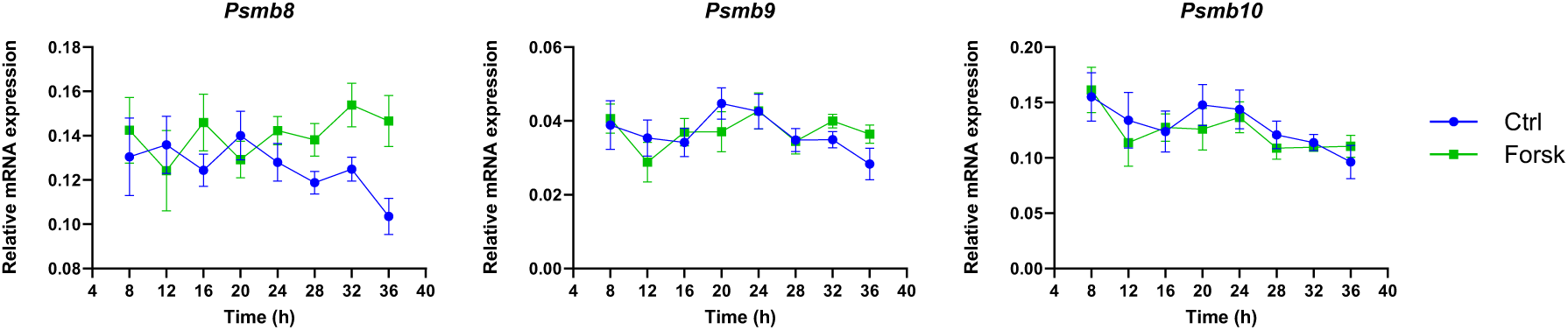
Forskolin pulsing does not result in rhythmic expression of inducible proteasome subunit genes. INS-1 cells were handled as described in FIG. S3. Relative mRNA expression is calculated using *Hprt1* and 5s rRNA as reference genes. Data are presented as means ± SEM (N=5-6).

**Figure S6.**
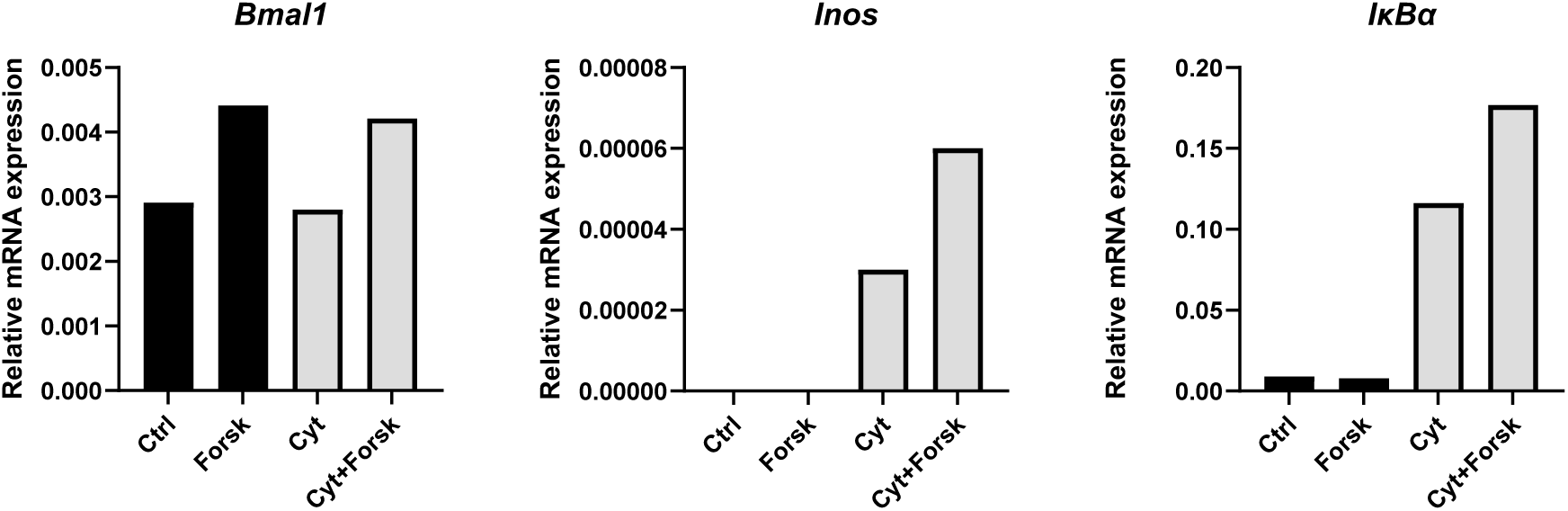
Synchronization potentiates the early cytokine-mediated NF-κB response. INS-1 cells were handled as described in FIG. 1, with the cells harvested after 4 hours of exposure. Relative mRNA expression was calculated using *Ppia* as a reference gene. Data are presented as means of 3 technical replicates (N=1).

**Figure S7.**
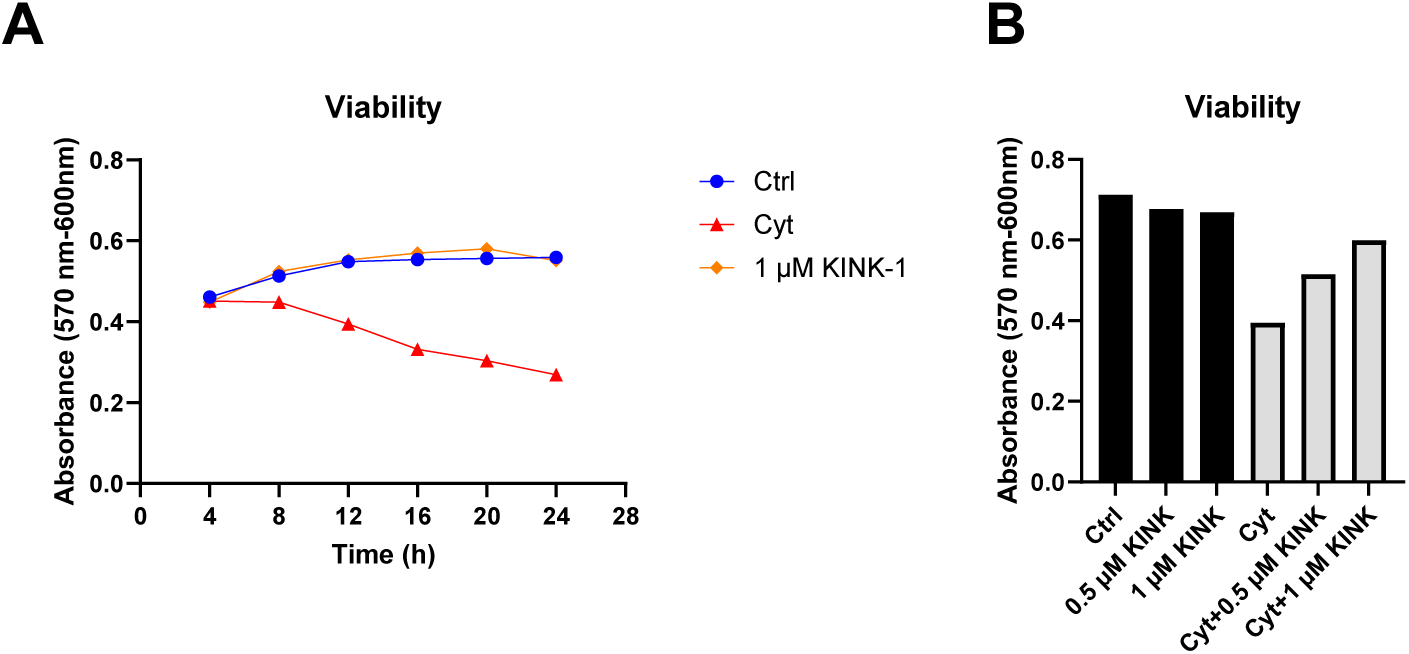
NF-κB inhibition attenuates cytokine-mediated alterations in clock gene expression, without affecting viability in non-cytokine exposed cells. INS-1 cells were cultured in the presence (Cyt) or absence (Ctrl) of 150 pg/ml of mIL-1β and 0.1 ng/ml rIFN-γ. Cells were pre-incubated with 0.5 or 1 µM KINK-1 (KINK) for one hour, and exposed to the inhibitor for the duration of the experiment. **(A)** Time-dependent changes in viability. **(B)** Dose-dependent effect of KINK-1 on viability. Data are presented as means of 3 technical replicates (N=1).

**Figure S8.**
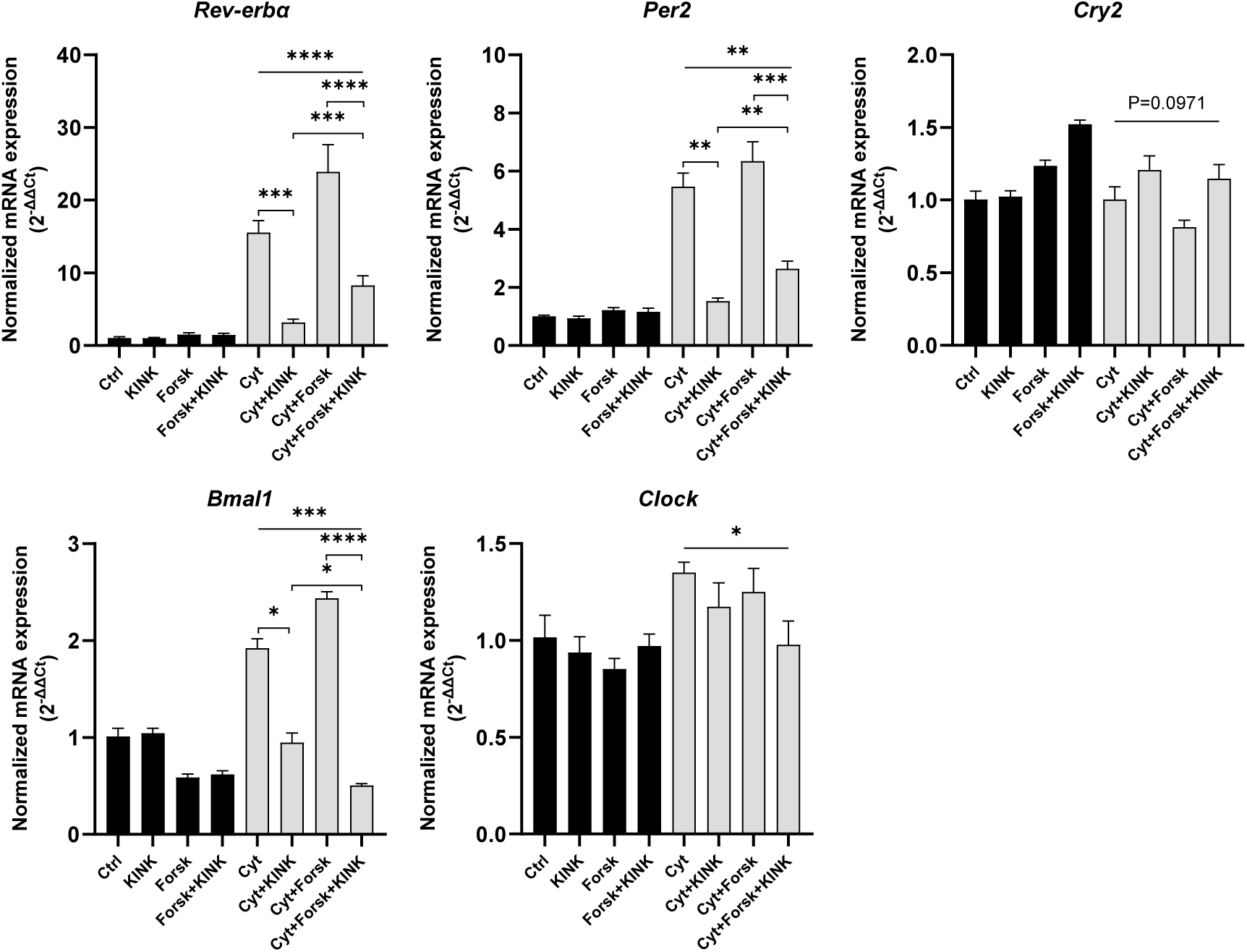
NF-κB inhibition differentially reverses proinflammatory cytokine modulation of core-clock gene expression. INS-1 cells were handled as described in FIG. 5C, and harvested after 20 hours. Normalized mRNA expression was calculated using *Hprt1* and 5s rRNA as reference genes. Data are presented as means ± SEM (N=4). Statistics are repeated measurements one-way ANOVA with p-values represented by symbols above the line and with Šídák’s corrected multiple comparisons tests.

**Figure S9.**
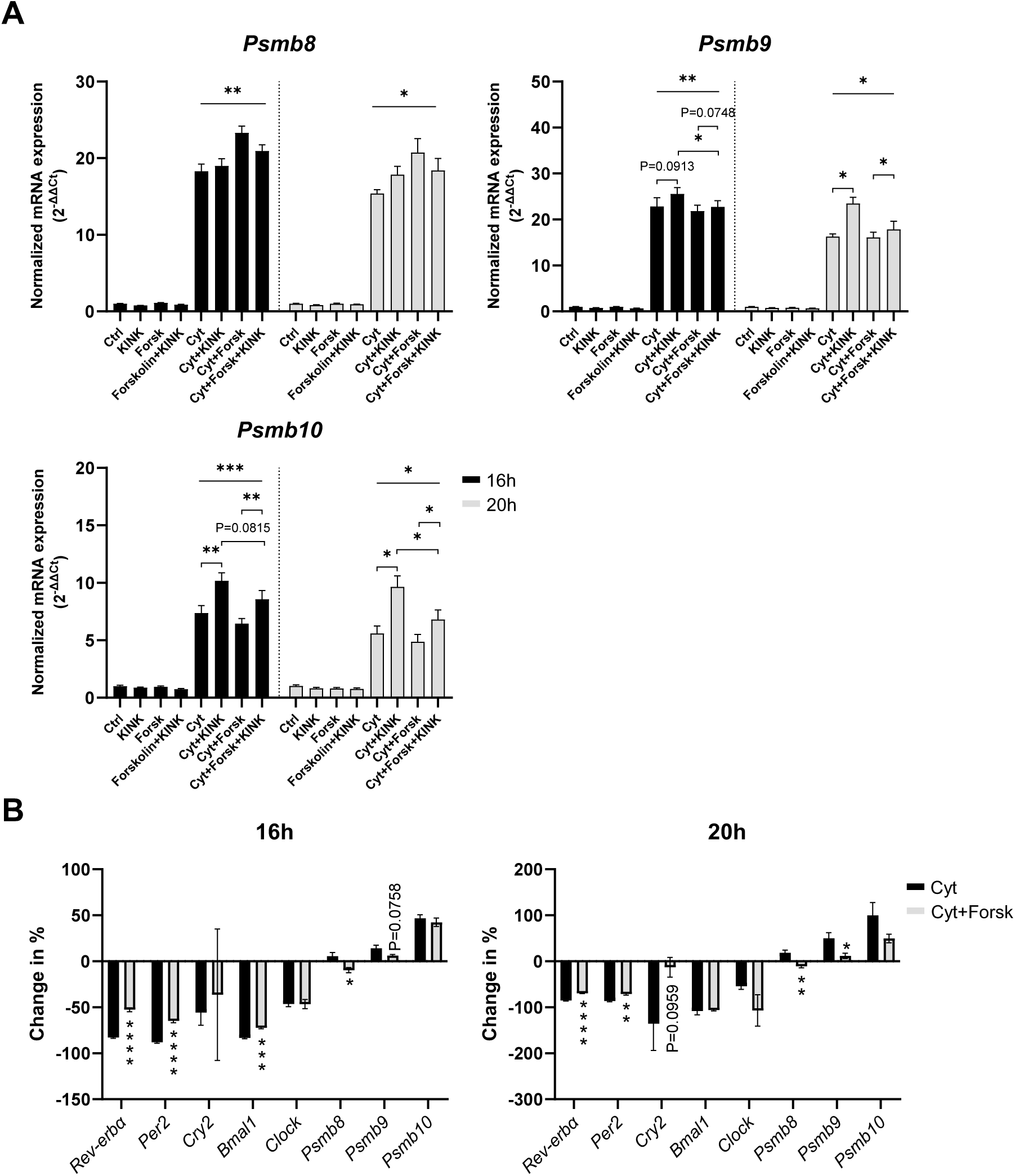
NF-κB inhibition differentially affects cytokine-mediated alterations in ind-proteasome subunits and core-clock gene expression in a synchronized system. **(A)** INS-1 cells were handled as described in FIG. 5C, and were exposed for either 16 or 20 hours. Normalized mRNA expression was calculated using *Hprt1* and 5s rRNA as reference genes. **(B)** Data from (A) and FIG. 5 presented as percentage change in expression after cytokine exposure caused by the addition of KINK-1, corrected for baseline expression level in the respective non-cytokine exposed controls. Data are presented as means ± SEM (N=4). Statistics is repeated measurements one-way ANOVA with p-values represented by symbols above the line and with Šídák’s corrected multiple comparisons tests (A), or two-tailed paired Student’s t-test (B).

**Figure S10.**
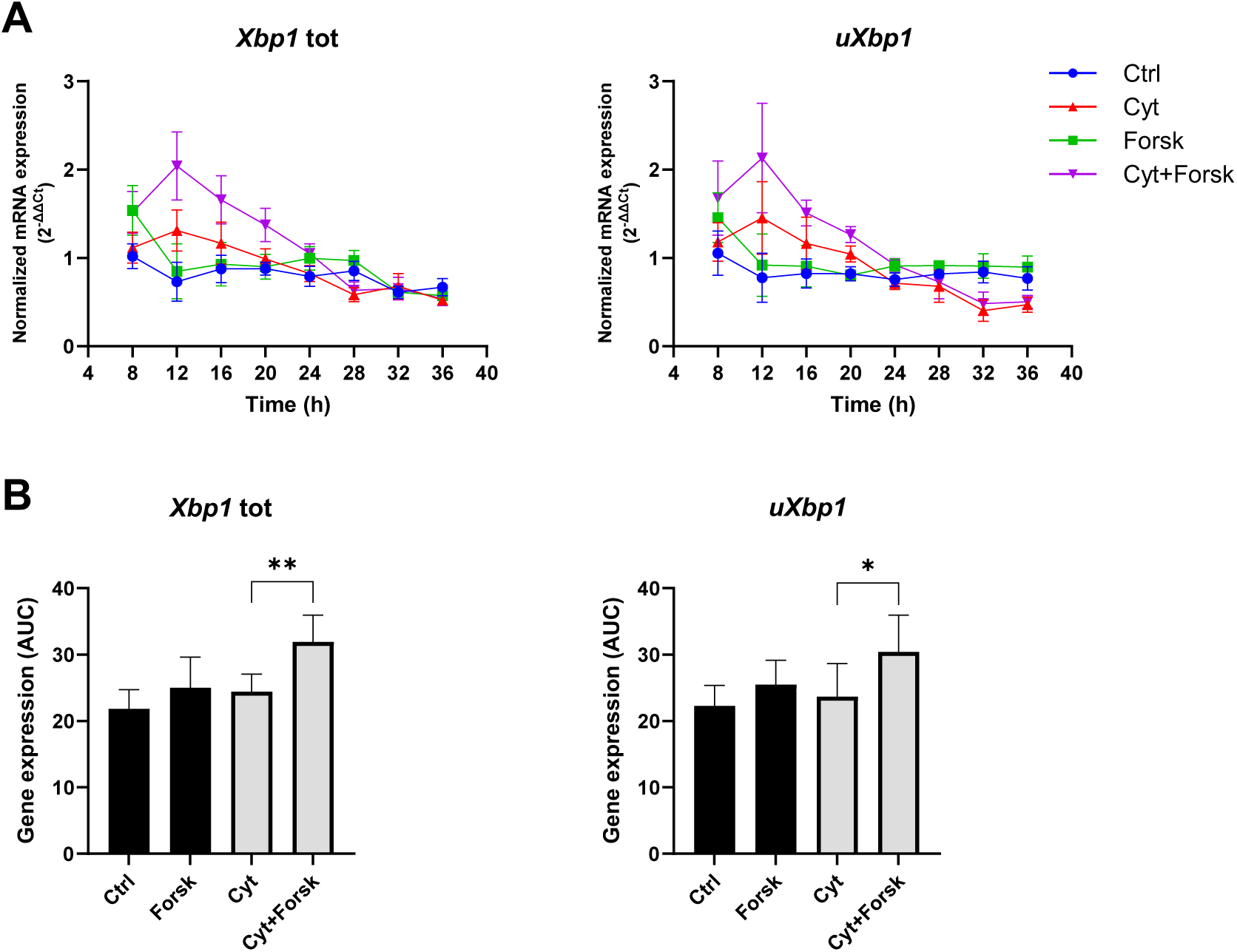
Synchronization augments cytokine-induced total *Xbp1* and u*Xbp1* expression. INS-1 cells were handled as described in FIG. 1. Normalized mRNA expression was calculated using *Hprt1* and 5s rRNA as reference genes. **(A)** Time-dependent normalized mRNA expression. **(B)** AUC of the curves in (A). Data are presented as means ± SEM (N=4). Statistics are two-tailed paired Student’s t-test.

**Figure S11.**
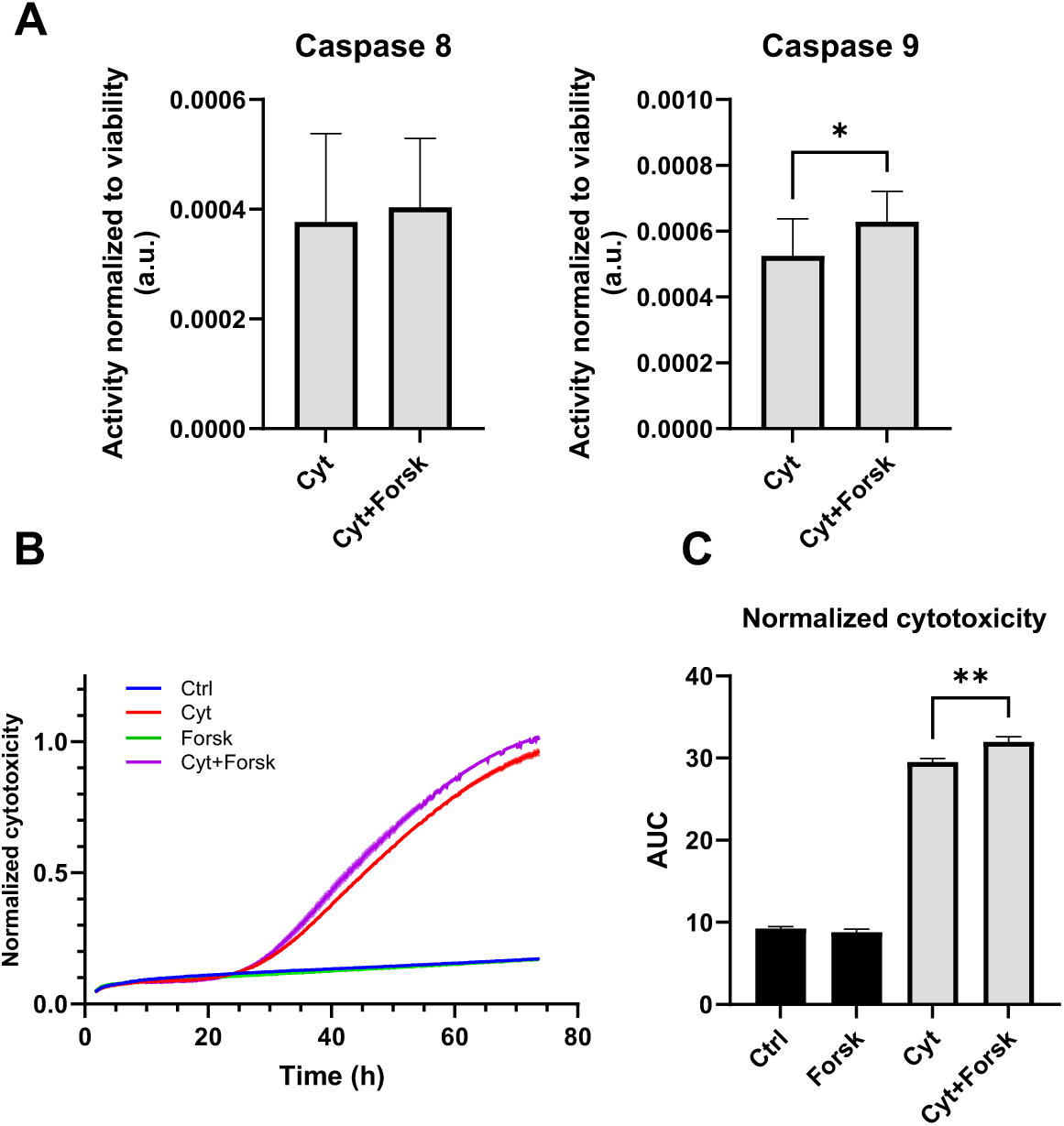
Synchronization potentiates cytokine-mediated cytotoxicity associated with an increase in Caspase 9 activity. INS-1 cells were handled as described in FIG. 2, without the interval harvesting. **(A)** Caspase activity normalized to viability after 24 hours of exposure. **(B)** Cytotoxic response normalized to the maximum signal obtained. Cytotoxicity was monitored in real-time for 72 hours at 10-minute intervals. **(C)** AUC of the normalized cytotoxicity response. Data are presented as means ± SEM (N=4). Statistics is two-tailed paired Student’s t-test.

**Figure S12.**
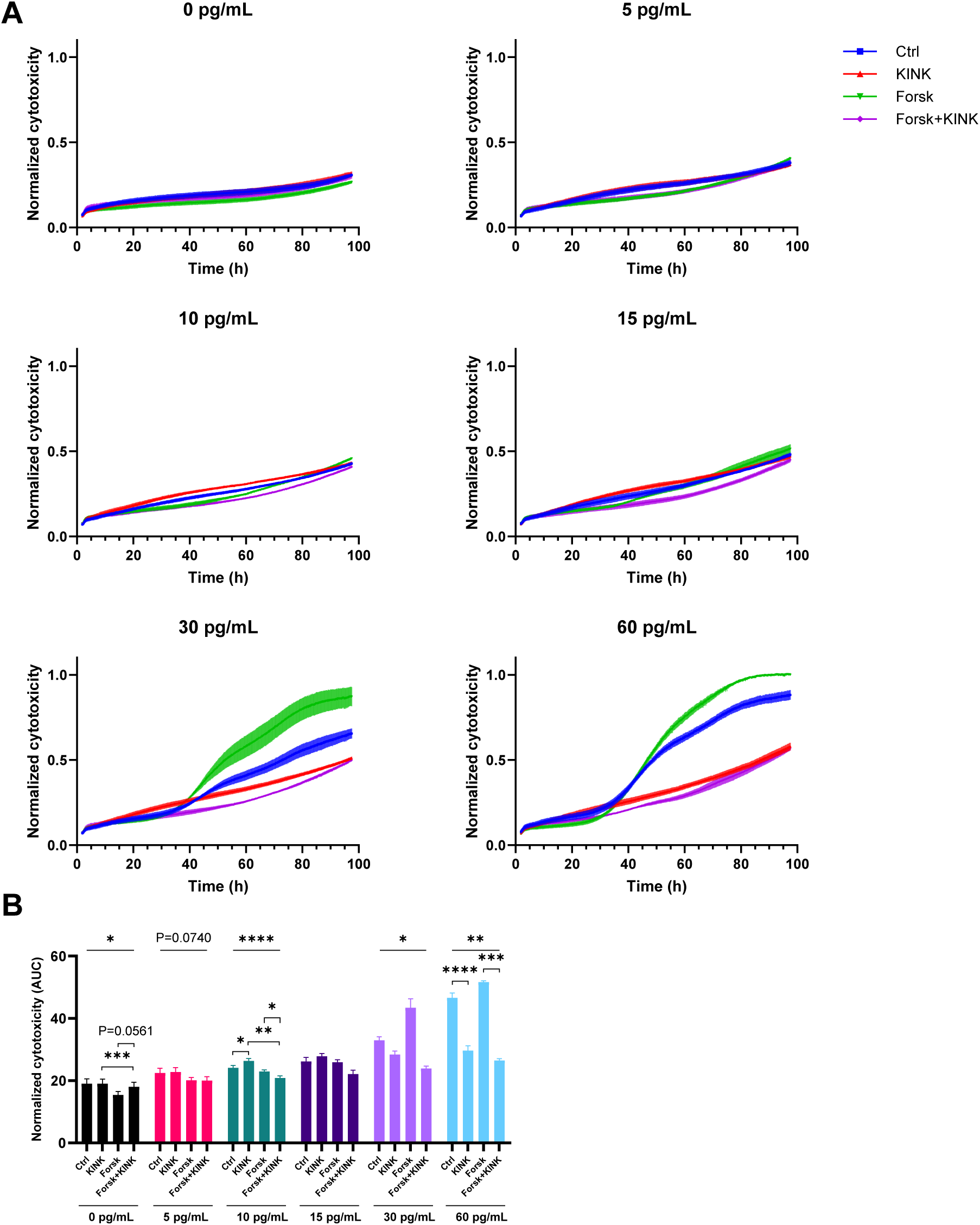
NF-κB inhibition alters cytotoxicity in synchronized INS-1 Per2 reporter cells depending on cytokine concentration. INS-1 Per2-luc cells were handled as described in FIG. 6. Annotated concentrations on plots are concentrations of mIL-1β+0.1 ng/ml rIFN-γ. **(A)** Cytotoxic response normalized to the maximum signal obtained. Cytotoxicity was monitored in real-time at 10-minute intervals for 96 hours, divided into single plots presenting different concentrations of mIL-1β. **(B)** AUC of the normalized cytotoxicity response. Data are presented as means ± SEM (N=3). Statistics is repeated measurements one-way ANOVA with p-values represented by symbols above the line and with Šídák’s corrected multiple comparisons tests.

**Figure S13.**
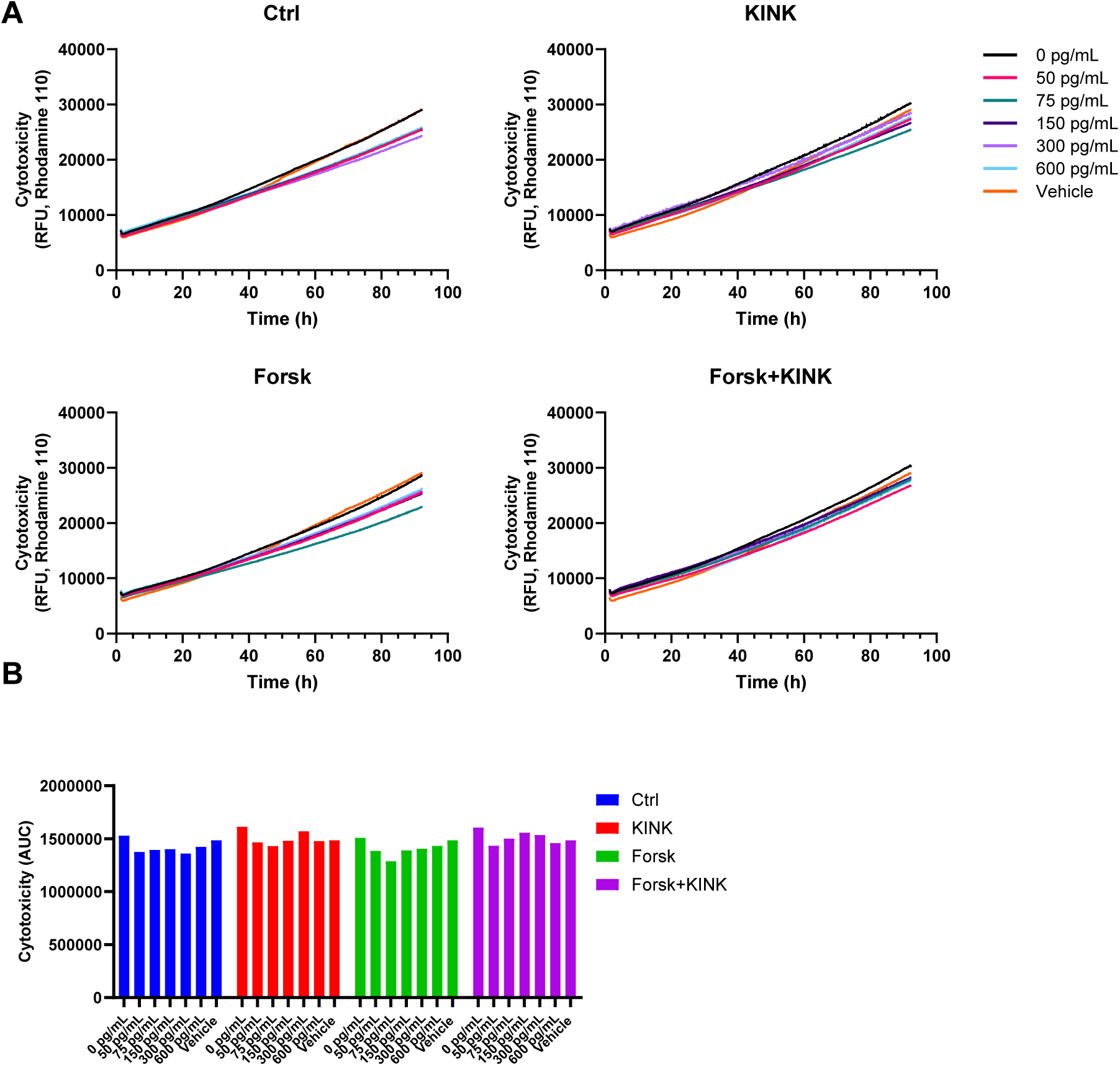
EndoC-βH3 cells do not respond to proinflammatory cytokines in a concentration-dependent manner. EndoC-βH3 cells were cultured in the presence or absence (ctrl) of varying concentrations of mIL-1β (legend) and 5 ng/ml hIFN-γ. In addition, cells were synchronized with a one-hour 10 μM forskolin pulse (Forsk), as well as 1 µM KINK-1 (KINK) in a one-hour preincubation and for the duration of the experiment. **(A)** time-dependent changes in cytotoxicity monitored in real-time at 10-minute intervals for 96 hours. **(B)** AUC of the cytotoxic response. Data are presented as means of three technical replicates (N=1).

**Figure S14.**
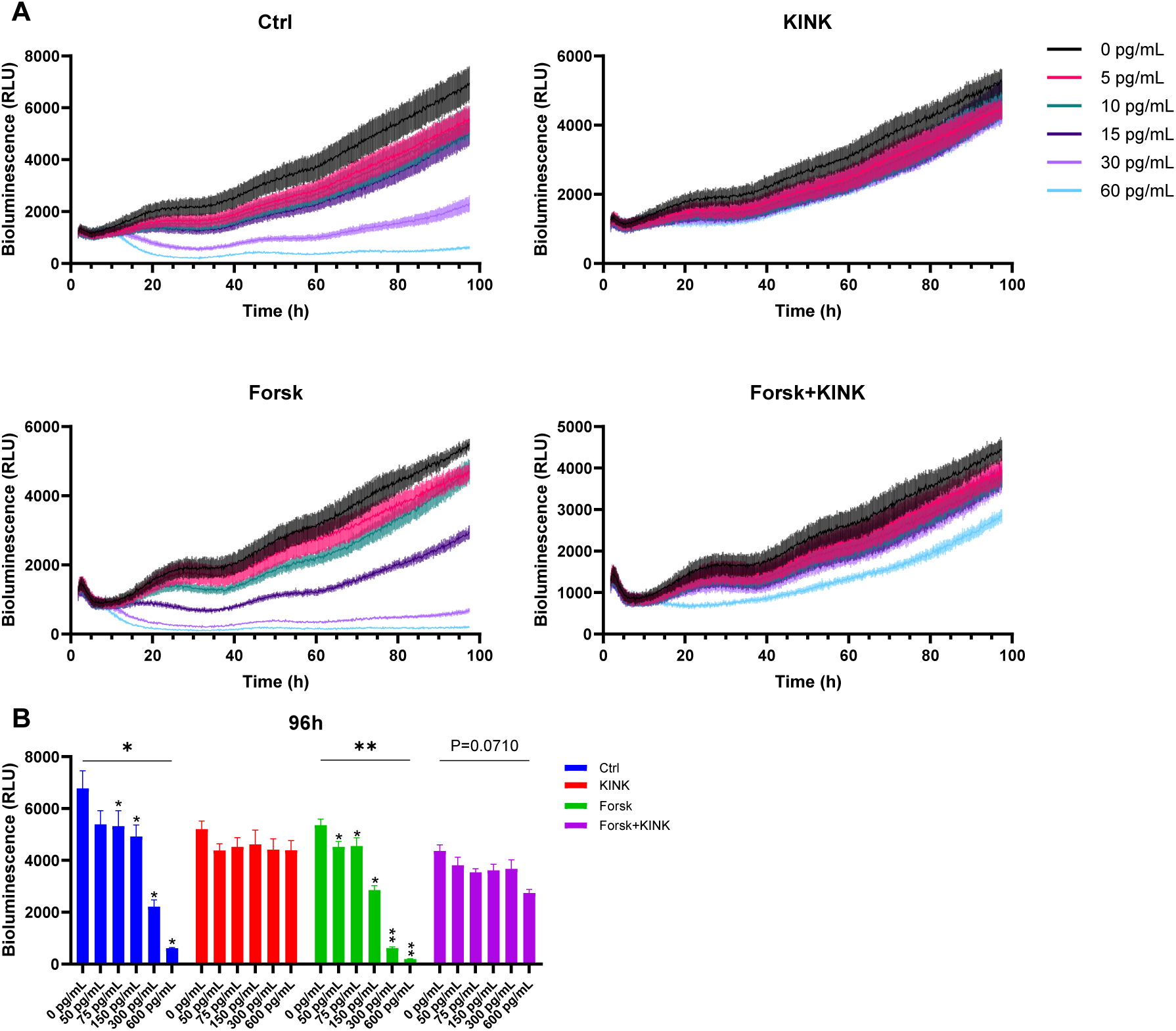
KINK-1 inhibits the proinflammatory cytokine-mediated reduction in bioluminescence. INS-1 Per2-luc cells were handled as described in FIG. 6. **(A)** Raw non-detrended bioluminescence traces of FIG. 7+8, recorded at 10-minute intervals for 96 hours. **(B)** Dose-response effect of proinflammatory cytokines on the 96-hour endpoint signal. Data are presented as means ± SEM (N=3). Statistics are repeated measurements one-way ANOVA with p-values represented by symbols above the line and with Dunnett’s corrected multiple comparisons tests.

**Figure S15.**
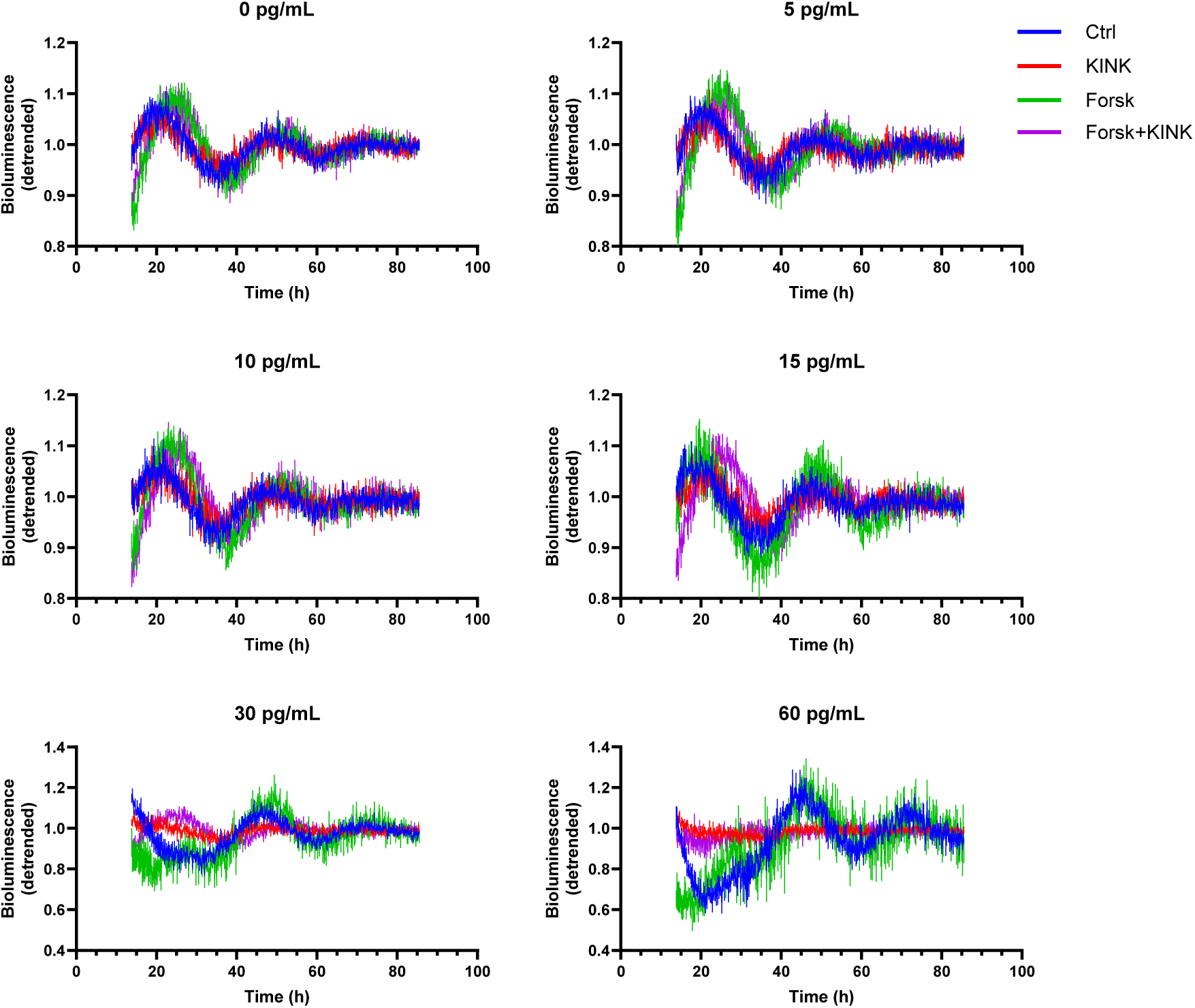
Concentration-dependent effects of proinflammatory cytokines on synchronized INS-1 Per2-Luc reporter oscillations. INS-1 Per2-Luc cells were handled as described in FIG. 6. **(A)** Detrended Per2-Luc bioluminescence signal presented in the functional window (13.5-85.5 hours) of the analysis, divided into single plots presenting different concentrations of mIL-1β+0.1 ng/ml IFN-γ. Bioluminescence was recorded at 10-minute intervals. Data are presented as means ± SEM (N=3).

**Figure S16.**
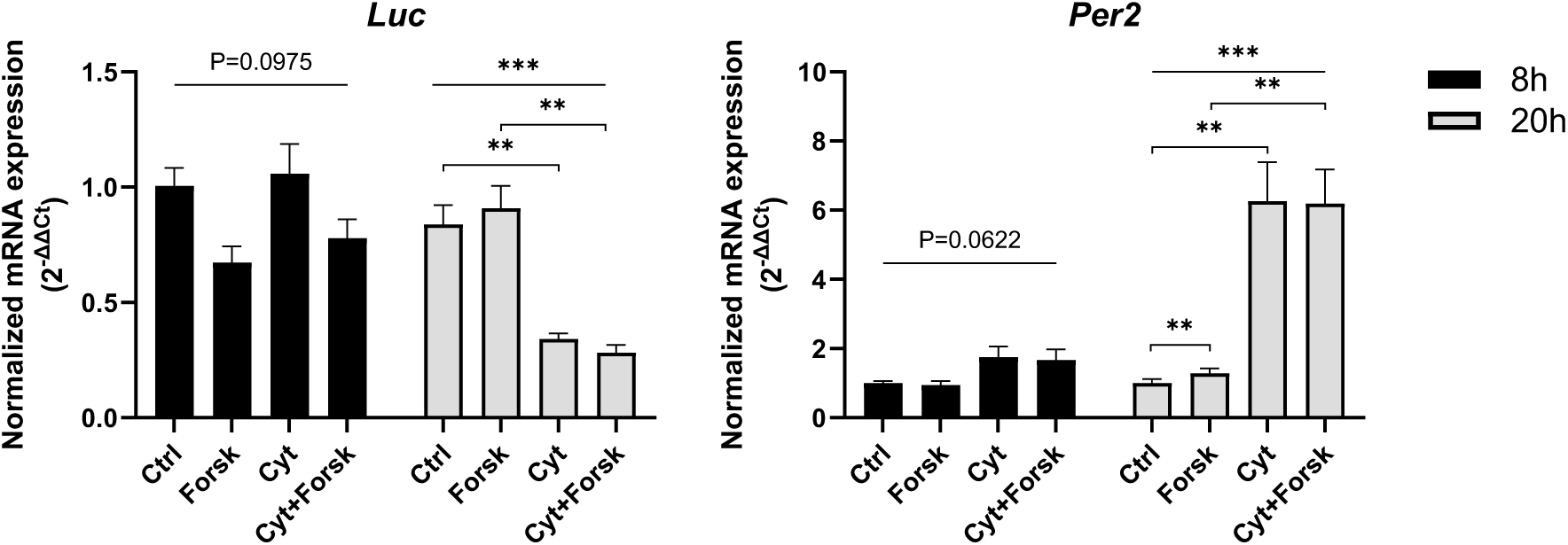
Cytokines reduce *Per2* promotor-driven *Luc* expression, due to increased repressive effect of *Per2*. INS-1 cells were handled as described in FIG. 1 and exposed for either 8 or 20 hours. Normalized mRNA expression is calculated using *Hprt1* and 5s rRNA as reference genes. Data are presented as means ± SEM (N=3). Statistics is repeated measurements one-way ANOVA with p-values represented by symbols above the line and with Šídák’s corrected multiple comparisons tests.

**Table S1.**
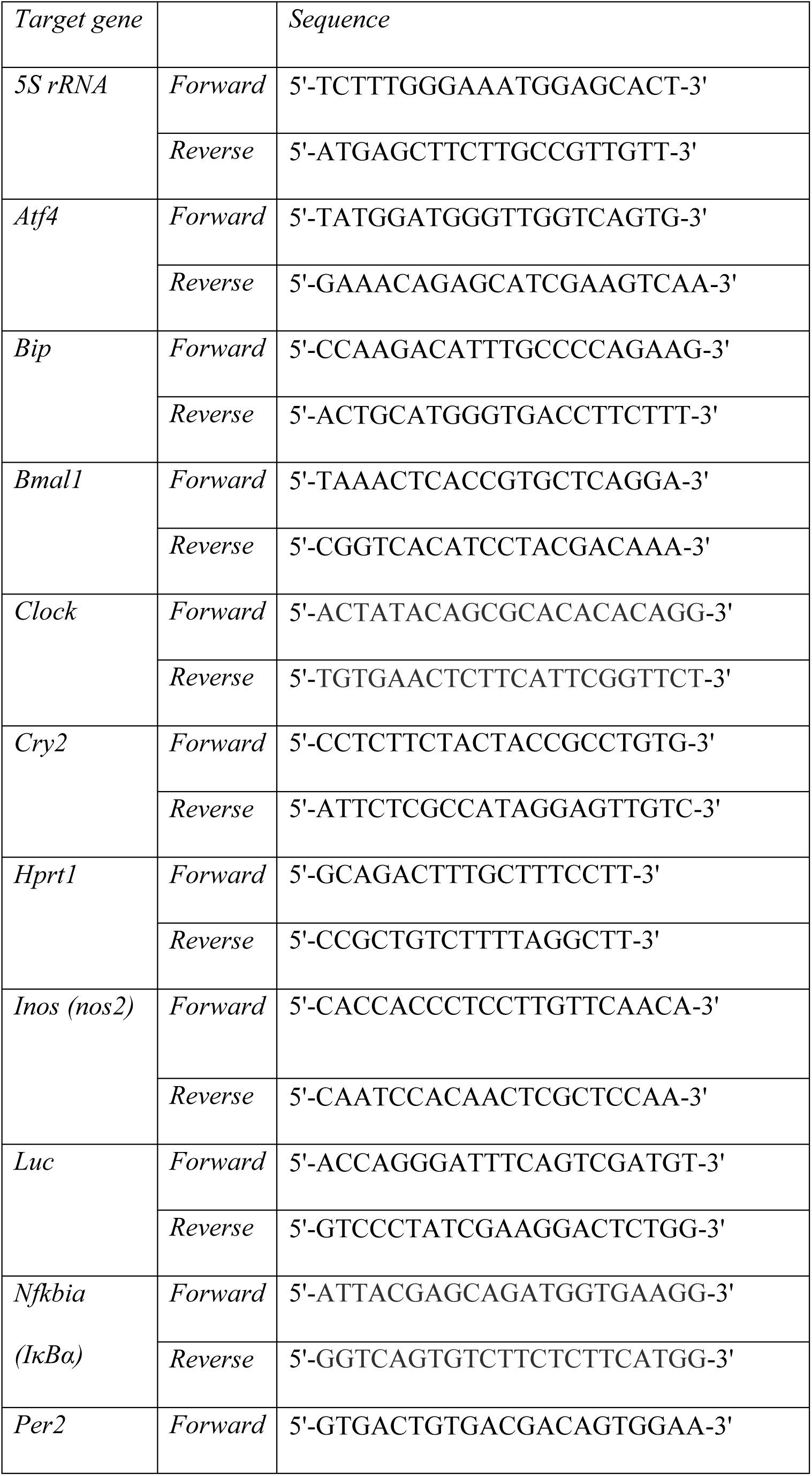
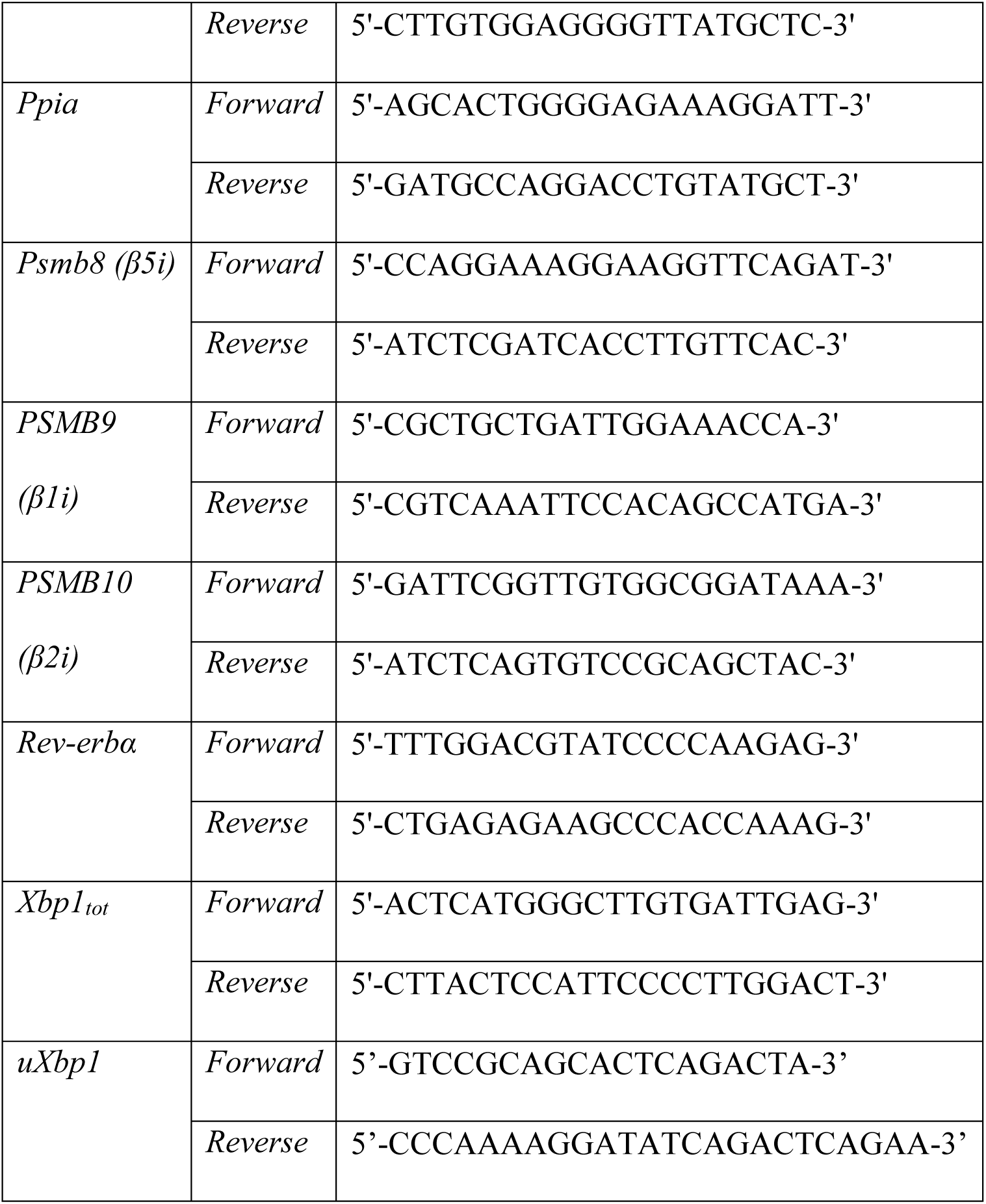
Gene-specific primers.

**Table S2.** Antibodies.

| **Antibody target** | **Company** | **Catalog number** | **Dilution** |
| --- | --- | --- | --- |
| Anti-mouse secondary | Cell Signaling | 7076P2 | 1:10 000 |
| Anti-rabbit secondary | Cell Signaling | 7074P2 | 1:10 000 |
| BMAL1 | Cell Signaling | D2L7G | 1:1000 |
| CLOCK | Santa Cruz | sc-271603 | 1:1000 |
| CRY2 | Invitrogen | PA5-79073 | 1:1000 |
| GAPDH | Abcam | ab8245 | 1:3000 |
| PER2 | MBL | PM083 | 1:500 |
| REV-ERBα | Cell Signaling | E1Y6D | 1:500 |

**Table S3.** Circadian rhythmicity of proteasome subunits. Table of proteasomal subunits and their rhythmic expression in islets. Cells marked in green display rhythmicity. Data from [36].

| *Gene name* | *Rhythmicity* |
| --- | --- |
| **20S Catalytic core particle (CP) – $\alpha$ subunits** |  |
| PSMA1 | Rhythmic |
| PSMA2 | Rhythmic |
| PSMA3 | Non-rhythmic |
| PSMA4 | Non-rhythmic |
| PSMA5 | Non-rhythmic |
| PSMA6 | Non-rhythmic |
| PSMA7 | Non-rhythmic |
| PSMA8 | N/A |
| **20S CP – $\beta$ subunits** |  |
| PSMB1 | Rhythmic |
| PSMB2 | Non-rhythmic |
| PSMB3 | Non-rhythmic |
| PSMB4 | Non-rhythmic |
| PSMB5 | Non-rhythmic |
| PSMB6 | Non-rhythmic |
| PSMB7 | Non-rhythmic |
| **20S CP – $\beta$ subunits (Inducible)** |  |
| PSMB8/ $\beta 5i$ | Rhythmic |
| PSMB9/ $\beta 1i$ | Rhythmic |
| PSMB10/ $\beta 2i$ | Rhythmic |
| **19S regulatory particles (RP) – ATPase subunits** |  |
| PSMC1 | Non-rhythmic |
| PSMC2 | Non-rhythmic |
| PSMC3 | Non-rhythmic |
| PSMC4 | Rhythmic |
| PSMC5 | Rhythmic |
| PSMC6 | Non-rhythmic |
| **19S RP – ATPase subunits – non-ATPase subunits** |  |
| PSMD1 | Non-rhythmic |
| PSMD2 | Rhythmic |
| PSMD3 | Rhythmic |
| PSMD4 | Non-rhythmic |
| PSMD6 | Non-rhythmic |
| PSMD7 | Non-rhythmic |
| PSMD8 | Non-rhythmic |
| PSMD11 | Rhythmic |
| PSMD12 | Non-rhythmic |
| PSMD13 | Rhythmic |
| PSMD14 | Non-rhythmic |
| ADRM1 | Rhythmic |
| SHFM1 | Non-rhythmic |
| **11S REG** |  |
| PSME1 | Rhythmic |
| PSME2 | Non-rhythmic |
| PSME3 | Non-rhythmic |
| **PA200** |  |
| PSME4 | Rhythmic |
| **PI31** |  |
| PSMF1 | Rhythmic in α-cells |

**Table S4.** Prediction of proteasomal and immunoproteasomal cleavage sites in clock proteins. Human and rat clock proteins (aa length in parenthesis) with the predicted number of proteasomal and immune proteasomal cleavage sites. The ratio is defined as “target sites”/”protein length”.

|  | Proteasome | Proteasome ratio | Immunoproteasome | Immunoproteasome ratio |
| --- | --- | --- | --- | --- |
| BMAL1 |  |  |  |  |
| BMAL1_HUMAN (626 aa) | 175 | 0.279 | 251 | 0.401 |
| BMAL1_RAT (626 aa) | 173 | 0.276 | 252 | 0.403 |
| CLOCK |  |  |  |  |
| CLOCK_HUMAN (846 aa) | 232 | 0.274 | 301 | 0.356 |
| CLOCK_RAT (862 aa) | 229 | 0.266 | 303 | 0.352 |
| CRY1 |  |  |  |  |
| CRY1_HUMAN (586 aa) | 191 | 0.325 | 261 | 0.445 |
| CRY1_RAT (588 aa) | 190 | 0.323 | 260 | 0.442 |
| CRY2 |  |  |  |  |
| CRY2_HUMAN (593 aa) | 193 | 0.325 | 266 | 0.449 |
| CRY2_RAT (594 aa) | 201 | 0.338 | 262 | 0.441 |
| PER1 |  |  |  |  |
| PER1_HUMAN (1290 aa) | 293 | 0.227 | 431 | 0.334 |
| PER1_RAT (1293 aa) | 297 | 0.229 | 434 | 0.336 |
| PER2 |  |  |  |  |
| PER2_HUMAN (1255 aa) | 317 | 0.253 | 457 | 0.364 |
| PER2_RAT (1257 aa) | 316 | 0.251 | 457 | 0.364 |
| REV-ERBα | | | | |
| NR1D1_HUMAN (614 aa) | 172 | 0.280 | 226 | 0.368 |
| NR1D1_RAT<br>(615 aa) | 174 | 0.283 | 228 | 0.371 |

